# Filling the Spinal Fracture Treatment Gap: An Osteoporotic Rabbit Model of Vertebral Augmentation

**DOI:** 10.64898/2025.12.18.695239

**Authors:** August J. Hemmerla, Abigail R. Grisolano, Austin D. Kimes, John T. Wray, Samantha E. Huddleston, Shwetha Ramachandra, Ryan E. Schultz, Don K. Moore, Ji-Hey Lim, Bret D. Ulery

## Abstract

**Background context:** Vertebral compression fractures (VCFs) are the most common osteoporotic fracture, yet current vertebral augmentation (VA) approaches rely exclusively on poly(methyl methacrylate) (PMMA), a non-resorbable cement that is associated with significant clinical drawbacks. Preclinical models are essential for evaluating novel injectable, bioactive alternatives, but existing small- and large-animal systems have translational limitations.

**Purpose:** The purpose of this study was to develop and validate a reproducible osteoporotic rabbit model that integrates controlled osteoporosis induction with a surgically accessible, fluoroscopically guided VA procedure suitable for preclinical evaluation of novel injectable biomaterials.

**Study design:** This study employed a basic science, preclinical method-development approach using an adult female New Zealand White rabbit model of osteoporosis combined with lumbar VA.

**Methods:** Adult female New Zealand White rabbits underwent bilateral ovariectomy followed by glucocorticoid treatment to establish osteoporosis as confirmed by qCT-derived reductions in bone mineral density and T-scores ≤ -2.5. A novel open dorsal surgical approach was optimized *ex vivo* and applied *in vivo* to access the lumbar vertebrae (L4 - L5) for bilateral cortical decortication and PMMA injection. Structural changes were assessed by quantitative CT before and after VA.

**Results:** The induction protocol produced consistent, reversible, and re-inducible osteoporotic phenotypes, with significant reductions in trabecular and cortical density as well as thinning of cortical microarchitecture. The surgical workflow achieved reproducible, fluoroscopically-confirmed cement delivery into the vertebral cancellous compartment while minimizing leakage. Post-VA imaging demonstrated stable trabecular and cortical parameters, with expected reductions in bone surface metrics attributable to surgical access and cement filling.

**Conclusion:** This study establishes the technical feasibility and reproducibility of a combined osteoporotic and VA rabbit model that integrates disease-relevant bone degeneration with a clinically aligned surgical workflow.

**Clinical significance:** This model bridges a critical translational gap between rodent and large-animal systems and provides a robust preclinical platform for evaluating next-generation injectable, bioactive, and biodegradable materials aimed at improving safety and regenerative outcomes in osteoporotic VCF repair.

## Introduction

Osteoporosis is a progressive systemic skeletal disorder characterized by low bone mineral density (BMD) as well as trabecular and cortical microarchitecture deterioration that all together significantly weakens overall bone integrity and strength. [1] The World Health Organization specifically defines osteoporosis as having a trabecular bone score (T-score) at least 2.5 standard deviations below healthy BMD, while scores 1.0 - 2.5 standard deviations below healthy levels indicate osteopenia (*i.e.*, pre-osteoporosis). [2] Bone loss typically begins to occur in patients in their late 30s to early 40s, starting at a gradual pace that accelerates in the decades that follow, [3] particularly in postmenopausal women, [4] resulting in approximately 10% of those over 50 years old in the United States suffering from osteoporosis. [5] Specifically, age-related changes in bone quality due to osteopenia or osteoporosis are a major contributor to fragility fractures, playing a crucial role in why roughly half of all postmenopausal women have broken a bone. [6, 7] In the United States, more than 3 million osteoporotic fractures occur each year which are responsible for $25.3 billion in annual associated costs. [8] Specifically, vertebral compression fractures (VCFs) are the most common osteoporosis-related fracture, accounting for about 27% of all breaks, followed by wrist and pelvic fractures at 19% and 14%, respectively. [9, 10] Clinically, VCFs frequently involve the thoracic and lumbar sections of the spine and are typically caused by low-energy trauma that leads to vertebral body collapse. [11] These fractures are often quite symptomatic, presenting with persistent pain, kyphotic deformity (*i.e.*, excessive spine curvature), reduced pulmonary function, and, in the most severe cases, neurological deficits. [12]

Conservative care for VCFs includes medication, bracing, physical therapy, and nerve root blocks, all of which offer varying degrees of pain relief and functional improvement. [13] When these measures fail to provide adequate relief or when persistent pain continues to significantly affect quality of life, surgical intervention through vertebral augmentation (VA) may be considered and is used in nearly one-fifth of all symptomatic VCFs cases. [14] The most common VA procedures — percutaneous vertebroplasty and percutaneous kyphoplasty — both involve the injection of poly(methyl methacrylate) (PMMA) bone cement into the fractured vertebral body. The primary difference between the two procedures is that the former technique involves just a PMMA injection whereas the latter approach attempts to restore vertebral body height using balloon inflation before the PMMA is delivered. Both vertebroplasty and kyphoplasty have been shown to be quite effective at reducing pain and improving mobility. [15, 16] Unfortunately, PMMA is the only currently FDA-approved material for use in these procedures, [17, 18] even though it has several significant drawbacks including cytotoxicity of its components, [19] tissue necrosis due to its high exothermic curing reaction, [20] mechanical properties that greatly exceed those of native cancellous bone, [21] and potential to leak into surrounding tissues. [22] The last of these problems is particularly concerning as it can lead to devastating complications such as inducing a pulmonary embolism or causing a spinal cord injury. [23] To overcome these material deficits, research has explored the incorporation of various fillers and functional additives to improve the bioactivity, dispersion, and structural integration of PMMA. [24, 25] Regardless, due to the non-resorbable nature and long-term persistence of PMMA, there still exists a need to develop injectable, bioactive, and biodegradable materials that provide transient mechanical support while promoting *in situ* bone regeneration.

Given the need for improved material design, animal models play a crucial role in the preclinical evaluation of new products for VA-based VCF repair. Among the currently available animal models, sheep VA is the most clinically relevant, as this species mimics human bone and soft tissue well while offering more similar spinal biomechanics to humans than other species. [26, 27] Unfortunately, inducing osteoporosis in large animals like sheep, as well as pigs and dogs, is much more complex and variable than with smaller animals, often requiring a combination of ovariectomy (OVX), low-calcium diet (LCD), and gluticorticoid (GC) treatment over quite prolonged periods of time (*i.e.*, 6 - 18 months) to mimic disease. [27–29] Additionally, skeletally mature large animals must be used, as younger individuals often fail to exhibit sufficient significant bone loss even when employing induction protocols. In many cases — particularly in dogs and minipigs — only mild osteopenia can be achieved, [30] so often bone regenerative materials are only studied in healthy large animal models. [26]

By contrast, small animal models offer greater experimental flexibility and can be made osteoporotic more reproducibly. Rodents have been frequently used in osteoporosis research due to their low cost, relatively short life cycle, and ease of handling, as well as their ability to present an osteoporotic phenotype by OVX often combined with LCD and/or GC treatment. [27, 31–33] However, the small size of rodent vertebrae, their rapid bone healing, and their continued spinal growth, limits their utility for evaluating the long-term performance of implanted VA materials. Additionally, anatomical differences — such as the absence of a fully developed Haversian system in their vertebrae — further reduce the relevance of the rat model in recapitulating human vertebral degeneration and remodeling. [34] In contrast, osteoporotic rabbit models offer several advantages as a translational model in osteoporosis research including their moderate cost, ease of handling, larger bone size, and the presence of a Haversian system that supports bone remodeling. [35] While some rabbit VA models have been described in the literature, they remain limited in depth especially for the induction of osteoporosis before VA is performed. [36–40] Therefore, there exists a need for a more comprehensive rabbit model to be developed that best serves as an intermediate platform between small rodent and large animal models, improving the clinical translation pathway for novel VA materials.

Specifically, our enclosed work focused on inducing osteoporosis in adult New Zealand White female rabbits using an OVX+GC protocol. This was followed by a novel VA surgical procedure first developed through iterative *ex vivo* experiments to ensure anatomical and procedural consistency followed by *in vivo* refinement. The final protocol generated consists of an open dorsal approach with bilateral cortical defects created in the L4 and L5 vertebra to access the centrum’s cancellous region followed by PMMA injection to demonstrate the feasibility and reproducibility of this intersectional osteoporotic and VA surgical model. Vertebral microarchitectural parameters were assessed pre-VA and post-VA to characterize bone structural changes. This study establishes the technical feasibility of performing VA reliably in an osteoporotic rabbit model, providing an exciting foundation for future materials evaluation.

## Materials and Methods

### Animal Model and Experimental Design

Twenty-four adult female New Zealand White rabbits (2.7 - 3.0 kg) were obtained from Charles River. Animals were housed individually in an AAALAC-accredited vivarium [41] maintained at 20 - 22 °C on a 12-hour light/dark cycle, with *ad libitum* access to water and standard rabbit pellets. All animal procedures were approved by the University‘s Institutional Animal Care and Use Committee (IACUC Protocol #34201) and directly adhered to best practice guidelines for humane animal use.

### Ovariectomy (OVX) Procedure

To generate a desirable osteoporotic rabbit VA model, all 24 rabbits received an OVX followed by GC treatment before a VA was performed. Following a one-week acclimation period, all animals underwent a bilateral OVX after which daily health monitoring was conducted throughout the entire study period (). Under aseptic conditions, animals were given preemptive buprenorphine SR (0.1 mg/kg subcutaneously - SQ) and general anesthesia was induced with ketamine (10 mg/kg intramuscularly - IM) and dexmedetomidine (0.025 mg/kg IM). Anesthesia was further induced with 3 - 4% isoflurane delivered via a non-rebreathing circuit which was then maintained at 2 - 3.5%. Anesthetic depth was monitored by assessing vital signs and mucous membrane color at 15-minute intervals. Ophthalmic ointment was used to protect animal corneas and body temperature was maintained by warm water blankets and heat lamps as anesthetic central nervous system depression can impair thermoregulation. A 4 - 5 cm ventral midline incision was made caudal to the umbilical scar to expose the linea alba, which was dissected along the midline to expose the cecum and uterine horns. The uterine horns were retracted and exteriorized, after which the ovaries were ligated using 3-0 absorbable Securosorb™ suture. After organ removal, muscle and subcutaneous tissue were closed using 3-0 absorbable suture and the skin was apposed with 4-0 absorbable Monocryl™ suture using a buried knot technique. Local analgesia consisting of bupivacaine (1 mg/kg) and/or lidocaine (1 mg/kg) were applied for post-operative incision pain, after which animals were monitored daily for 7 - 10 days as they recovered.

### Osteoporosis Induction

One-week post-OVX, osteoporosis was induced by daily subcutaneous injections of 0.6 mg/kg dexamethasone sodium phosphate (GC) from weeks 1 through 11 (**Figure 1**). [35] A steroid taper was then employed from weeks 11 through 13 post OVX, which corresponds to approximately 14 to 16 weeks before VA was performed (**Figure 1**).

**Figure 1.**
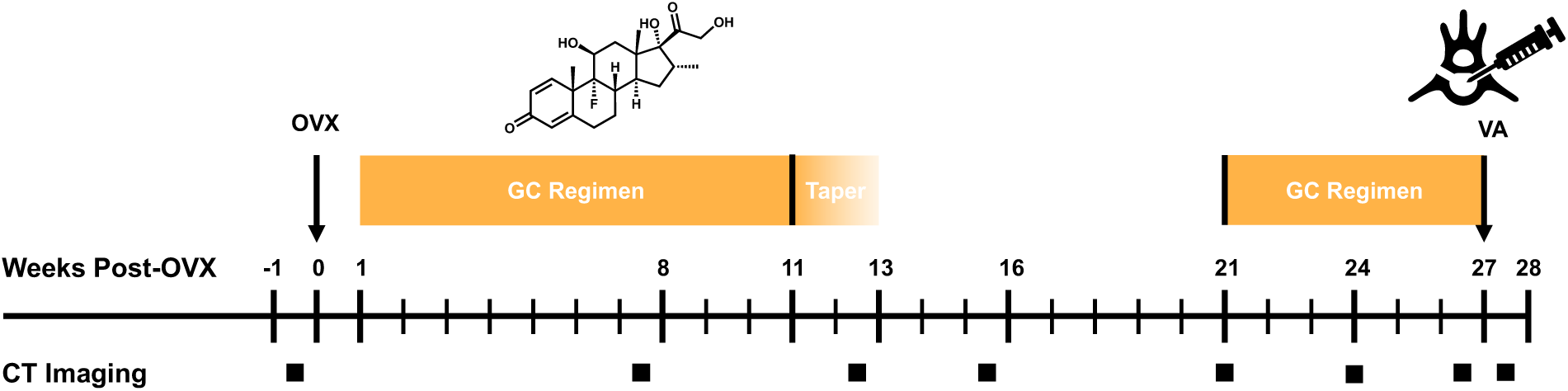
New Zealand White Rabbit OVX/VA Model Timeline. After a one-week acclimation period, rabbits underwent bilateral ovariectomy (OVX) followed by glucocorticosteroid (GC) administration to induce osteoporosis. Once sufficient bone loss was confirmed vertebroplasty (VA) was performed. Computed tomography (CT) scans were taken at noted intervals to monitor vertebral changes and treatment effects throughout the study.

### Quantitative Computer Tomography (qCT) Imaging and Analysis

qCT imaging was performed at specified timepoints (**Figure 1**) using a Siemens SOMATOM X.ceed 128-slice scanner. Helical acquisition parameters included 193 mAs reference, 120 kV tube voltage, 0.8 pitch, 1.0-second gantry rotation, and 0.5-mm slice thickness. Image reconstruction used the BR62 kernel and 0.5-mm increments. Multiplanar 3D reconstructions (*i.e.*, coronal, sagittal, and axial) were used to optimize visualization. BMD calibration was performed using a Mindways QCT Pro phantom scanned during a master calibration session. Rabbits were anesthetized for CT imaging using acepromazine (1 mg/kg SQ) followed by inhaled isoflurane (2 - 3%) supplemented with oxygen delivered via facemask. Animals remained under anesthesia for the duration of the imaging session and were monitored until fully recovered and ambulatory. Image analysis was performed using Dragonfly 3D World (Zeiss Edition 2024.1). Individual lumbar vertebrae (L1 – L7) segmentation was achieved through a combination of thresholding, connected components analysis, region-of-interest (ROI) selection, and watershed transformation. Cortical and trabecular regions in the L3 - L6 vertebrae were segmented using methods established by Kohler and colleagues (**Figure S1**) [42] with bone morphometric parameters computed according to Bouxsein and colleagues. [43] A full list of the quantitative measurements that were computed are provided in **Table S1**. PMMA was isolated from qCT images using threshold-based segmentation. Within each lumbar vertebra (*i.e.*, L4 and L5), PMMA ROIs were defined using upper and lower thresholds of ±3σ from the mean attenuation value. PMMA density was set at 1931.11 ± 15.05 mgHA·cm⁻³ (mean ± SD) and normality was confirmed using the Shapiro–Wilk test.

### Vertebral Augmentation (VA) Procedure

VA was conducted at 27 weeks post-OVX (**Figure 1**) on 8 rabbits employing the same anesthesia protocol as described for performing OVX. Rabbits were positioned in ventral recumbency on a radiolucent table. The lumbosacral region was shaved and prepped by alternating applications of chlorhexidine and 70% isopropyl alcohol. L4 - L5 pedicle identification was guided by manual palpation. As shown in **Figure 2**, a midline incision started at L7 exposed subcutaneous tissue and epaxial musculature. Parallel fascial incisions between the multifidus and longissimus muscles provided access to the L4 - L6 articular processes. The deep lumbar fascia was next incised bilaterally, and paraspinal muscles were subperiosteally reflected to expose two adjacent lumbar vertebrae (*i.e.*, L4 - L5, **Figure 2A**).

**Figure 2.**
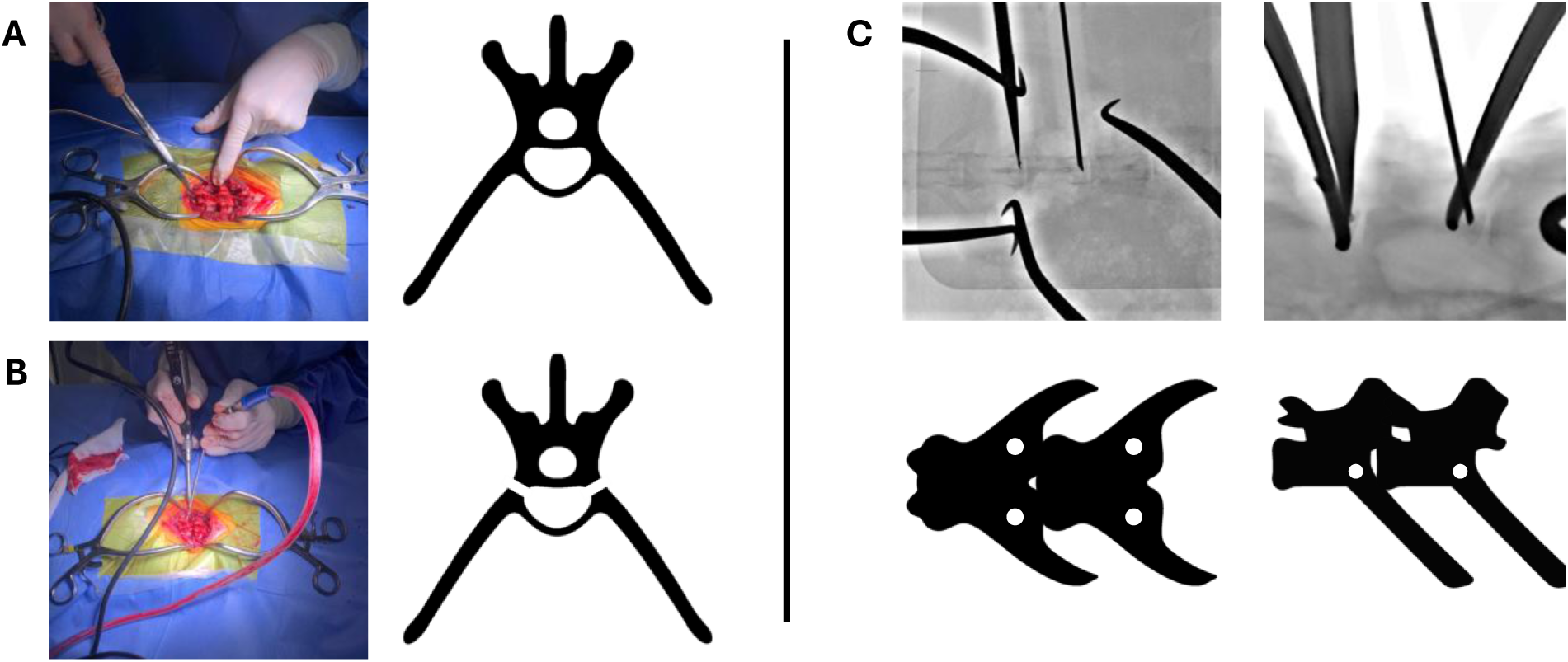
Operative alignment and imaging guidance for lumbar vertebral body access pre-injection. **(A)** A transverse schematic of the rabbit lumbar spine with corresponding intraoperative view demonstrates surgical exposure of L4 - L5. **(B)** A transverse schematic and paired intraoperative image illustrate bilateral decorticated portals created by burring through the dorsal cortex of each vertebral body. **(C)** Dorsal–ventral and sagittal schematics depict bilateral cortical defects at L4 and L5, with paired fluoroscopic images confirming accurate trajectory and access into the trabecular centrum.

An open, transpedicular approach was then used to create bilateral 1 mm decortication portals in both the L4 and L5 vertebra (**Figure 2B**) with a Stryker™ Core Sumex™ drill and 1 mm burr. Both the vertebral body and the injection site were confirmed fluoroscopically (Siemens Axiom Artis Cath/Angio, **Figure 2C**). Once situated, an estimated 0.05 - 0.1 mL of high-viscosity radiopaque PMMA cement (Kyphon HV-R, Medtronic), containing 30% BaSO₄ as a contrast agent, was injected per site via 16G needle under low pressure, with manual stabilization used to support polymerization (**Figure 3A**). Proper placement and final location of PMMA was visualized using fluoroscopy (**Figure 3B**). The full procedure duration ranged from 30 to 50 minutes (mean time of 42.7 ± 6.1 minutes) and postoperative monitoring mirrored that of the OVX protocol. Among the eight rabbits that underwent PMMA VA, immediate post-operative hind limb paralysis occurred in two animals, while delayed onset hind limb paralysis was observed in one animal at two weeks post-procedure. Complications were limited to the earliest procedures employing PMMA, whereas the final four surgeries resulted in the rabbits making a full normal recovery due to slowing the speed and reducing the quantity of PMMA injection to better suit the animal model used.

**Figure 3.**
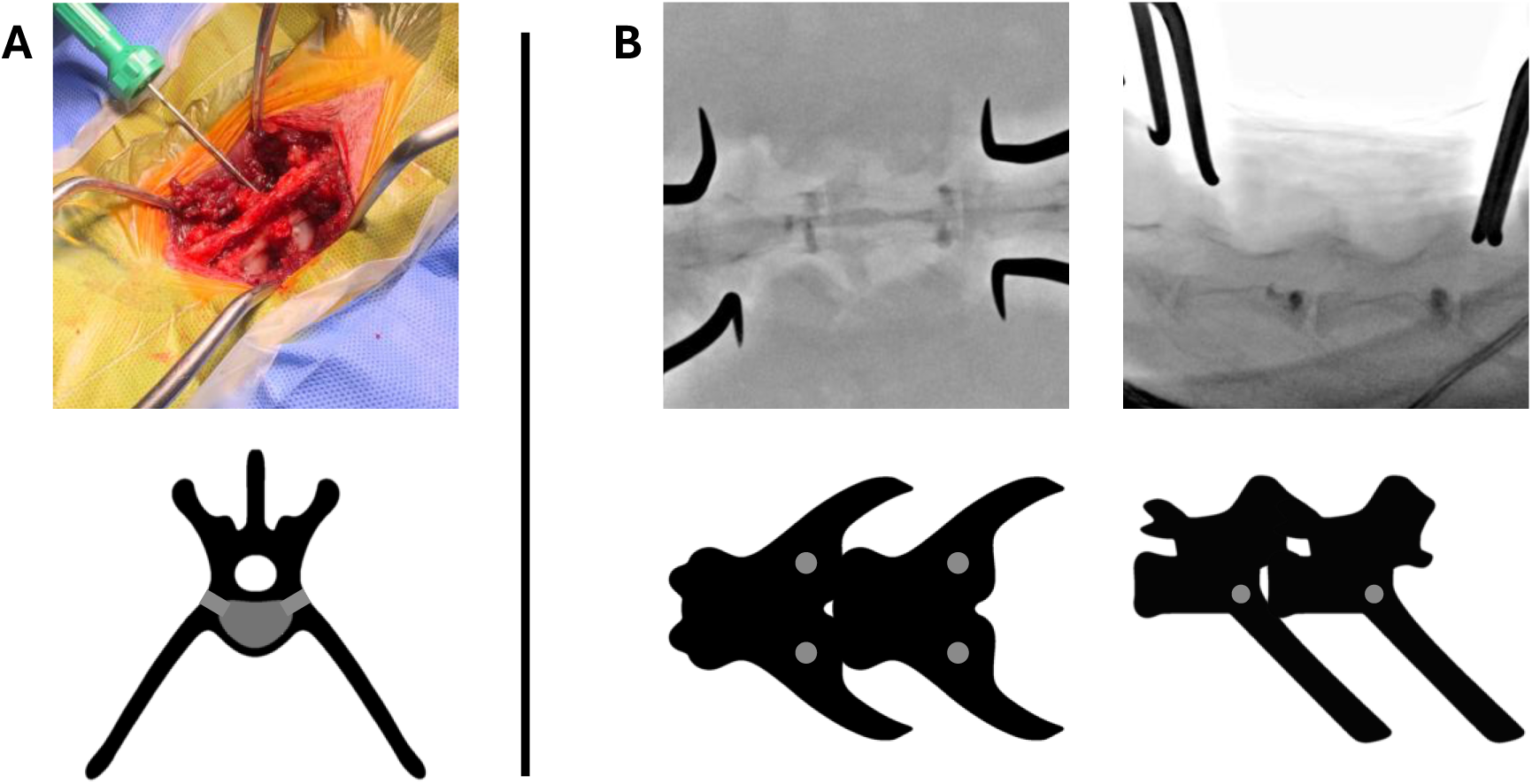
PMMA injection and fluoroscopic placement confirmation. **(A)** Surgical demonstration and schematic of VA injection. **(B)** Representative final fluoroscopic views show bilateral cement injection within the L4 and L5 vertebral bodies, confirming reproducible PMMA placement within the cancellous region.

### Exclusion Criteria & Euthanasia

All of the procedures above included exclusion criteria of intraoperative mortality, non-survival surgical outcomes, or postoperative hind limb paresis or paralysis. Regardless of study exclusion or inclusion, euthanasia for all New Zealand White rabbits included first anesthetizing with ketamine (25 mg/kg – IM) and xylazine (5 mg/kg – IM). Following adequate anesthesia, euthanasia was performed via intravenous injection of Euthasol (0.22 mL/kg of 86 mg/kg pentobarbital) into the ear vein with cervical dislocation used as a confirmatory method.

### Statistical Analysis

Statistical analysis was performed using JMP version 18 software. Data normality was first tested using the Shapiro-Wilk test. Hierarchical clustering was performed to evaluate similarity among vertebral levels based on normalized whole-bone, trabecular, and cortical qCT parameters. Clustering was implemented using Ward’s minimum variance method, which reduces the total within-cluster variance at each step of the agglomeration process. This analysis generated a dendrogram and a heatmap to visualize parameter relationships across lumbar levels (L3 - L6). Pooled and non-pooled group comparisons were then made using Analysis of Variance (ANOVA) with Tukey’s Honest Significant Differences (HSD) *post hoc* testing. Data are presented as mean ± standard deviation (N ≥ 5), for which groups that possess different letters have statistically significant difference in mean (p ≤ 0.05) whereas those that possess the same letter have similar means (p > 0.05).

## Results & Discussion

The model development timeline can be partitioned into three phases - ovariectomy, osteoporosis induction, and vertebral augmentation (**Figure 1**). Interestingly, before any procedure or protocol was undertaken (*i.e.*, at baseline week 0), healthy New Zealand White rabbits were found to possess measurable differences across their L3 - L6 lumbar levels for some parameters of interest (**Table S2**). While whole bone tissue mineral density (TMD.BV, 610.7 - 647.5 mgHA·cm⁻³), trabecular mineral density (TMD.Tb, 492.6 - 536.0 mgHA·cm⁻³), cortical mineral density (TMD.Co, 616.5 - 674.3 mgHA·cm⁻³), trabecular thickness (Tb.Th, 1.50 - 1.55 mm), and cortical thickness (Ct.Th, 1.16 - 1.33 mm) were statistically indistinguishable across vertebrae, volumetric measures showed significant differences. Specifically, total volume (TV) was reduced in L3 (2043 ± 171 mm³) when compared to L5 (2406 ± 119 mm³) and L6 (2388 ± 184 mm³). Also, bone volume (BV) was also lower in L3 (1877 ± 153 mm³) relative to L4 - L6 (2072 ± 102 mm³, 2236 ± 105 mm³, and 2245 ± 166 mm³, respectively) and the BV/TV ratio was decreased in L3 (0.919 ± 0.008) when compared to L6 (0.940 ± 0.012). Taken together, these findings indicate that L3 may be structurally distinct from L4 - L6, necessitating further analysis before lumbar data could be pooled across all multiple vertebrae. Therefore, hierarchical clustering of normalized whole-bone, trabecular, and cortical parameters was performed using Ward’s method (**Figure 4**), which groups vertebrae by minimizing differences in bone parameters across clusters. Heatmap analysis revealed greater variability across L3 - L6 for whole-bone and cortical parameters as compared to trabecular parameters. The dendrogram determined the L4 and L5 were most similar, followed by L6, with L3 being the most distinct from the other three. Based on this result, subsequent pooled analysis consisted of L4 - L6.

**Figure 4.**
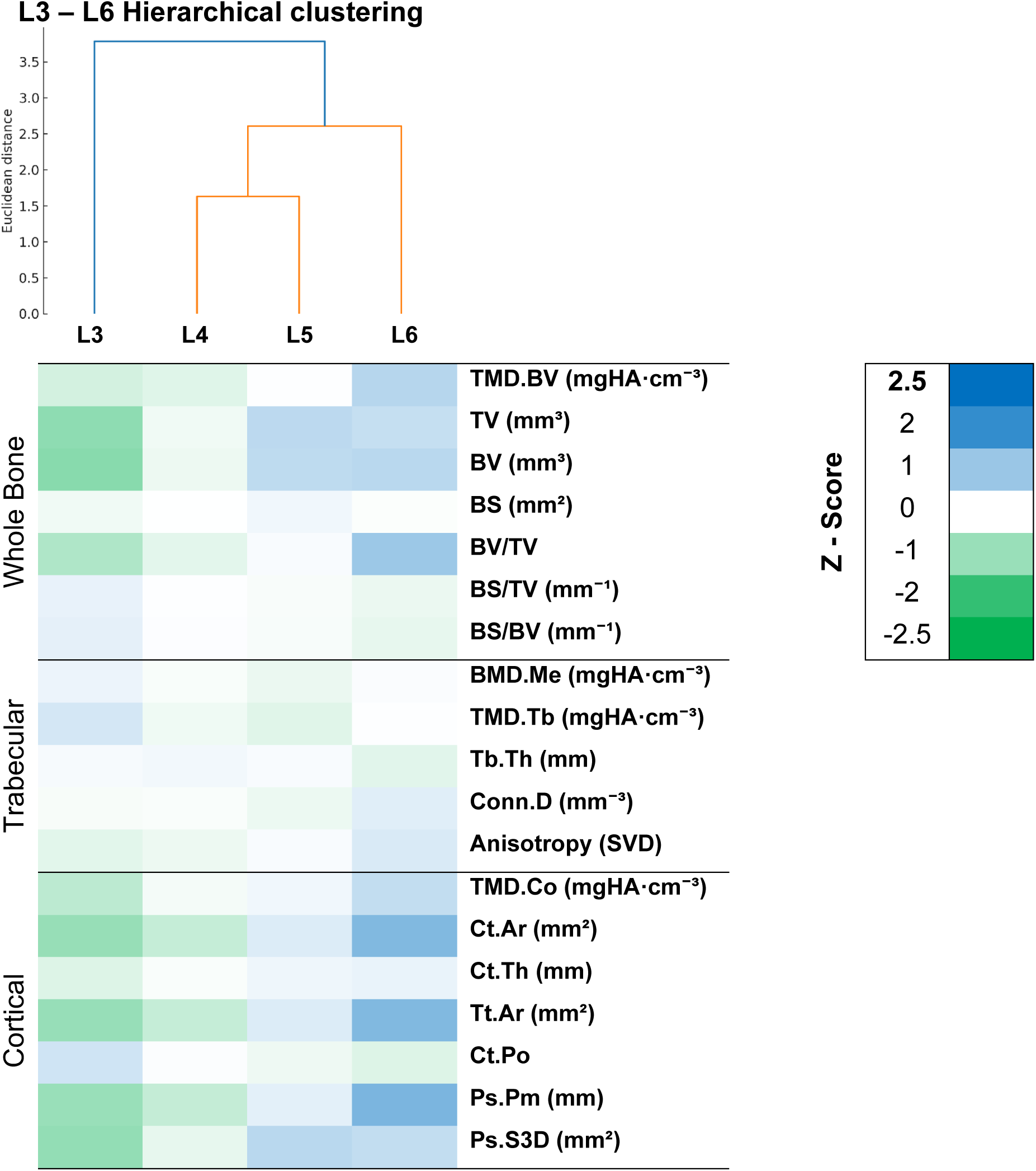
Summary of Healthy Rabbit Lumbar Vertebrae qCT Data. A heatmap of whole, trabecular, and cortical bone morphometric parameters (*i.e.*, rows) across lumbar vertebrae L3 - L6 at Week 0, representing healthy controls prior to any treatment, are provided. Values were z-scored within each variable to normalize differences in the measurement scales. The accompanying hierarchical clustering dendrogram (Euclidean distance, Ward’s method) showed L3 followed by L6 being more distinctly different than L4 and L5.

Following ovariectomy, 1 week of rest, and 7 weeks of glucocorticoid treatment, significant changes in some bone properties were observed (**Table S3**). Most interestingly, TMD.BV decreased by 23.8% from 629.1 ± 31.9 mgHA·cm⁻³ to 479.2 ± 38.5 mgHA·cm⁻³. Additionally, TMD.Tb declined by 24.2% (from 502.0 ± 52.1 mgHA·cm⁻³ to 380.6 ± 78.7 mgHA·cm⁻³), TMD.Co dropped by 26.3% (from 656.5 ± 37.7 mgHA·cm⁻³ to 484.1 ± 56.1 mgHA·cm⁻³), and BMD.Me was reduced by 26.6% (from 357.5 ± 79.9 mgHA·cm⁻³ to 262.4 ± 85.9 mgHA·cm⁻³). In contrast, TV (2344 ± 156 mm³ to 2411 ± 228 mm³) and BV (2184 ± 147 mm³ to 2203 ± 224 mm³) did not differ significantly, confirming that the initial osteoporotic state was driven by reductions in density rather than gross morphometry.

During the sustain/taper treatment phase (*i.e.*, weeks 8 to 13) as well as the drug-free interval (*i.e.*, weeks 13 to 21), partial recovery of bone properties was evident (**Table S4**). TMD.BV increased from 479.2 ± 38.5 mgHA·cm⁻³ to 609.2 ± 26.8 mgHA·cm⁻³, approaching the baseline value (*i.e.*, 629.1 ± 31.9). BMD.Me rose from 262.4 ± 85.9 mgHA·cm⁻³ to 268.1 ± 52.8 mgHA·cm⁻³, while BV recovered from 2203 ± 224 mm³ to 2396 ± 131 mm³, and TV jumped from 2411 ± 228 mm³ to 2595 ± 144 mm³. Despite these improvements, complete recovery was not achieved. Trabecular thickness (Tb.Th) declined modestly by 9.35% (from 1.573 ± 0.266 mm to 1.426 ± 0.115 mm) and cortical thickness (Ct.Th) rose only slightly by 13.3% (from 1.289 ± 0.533 mm to 1.461 ± 0.197 mm), suggesting that osteoporosis-induced microarchitectural alterations were only partially reversed.

Excitingly, a second GC regimen from week 21 to week 27 (**Figure 5A**) was able to re-establish an osteoporotic phenotype (**Figure 5B-D**). These structural deteriorations were qualitatively observed in representative CT images (**Figure 5B** and **Figure S2**), which showed pronounced trabecular rarefaction in the medullary cavity and cortical thinning along the articular and transverse processes. TMD.BV decreased significantly by 10.1% from 609.2 ± 26.8 mgHA·cm⁻³ to 547.7 ± 37.6 mgHA·cm⁻³ (**Figure 5C** and **Table S5**). The BV/TV ratio also fell from 0.924 ± 0.010 to 0.901 ± 0.023. Additional parameters that decreased include Tb.Th (from 1.426 ± 0.115 mm to 1.304 ± 0.150 mm) and Ct.Th (from 1.461 ± 0.197 mm to 1.116 ± 0.248 mm). Complementarily, cortical porosity (Ct.Po) jumped by 59.3% (from 0.172 ± 0.031 to 0.274 ± 0.091). At the start of the second GC regimen, rabbits possessed a modest decrease in bone density resulting from the GC taper, with T-scores of -0.62 ± 0.84 at week 21. With reinitiation of GC, the T-score fell over the next three weeks reaching -1.37 ± 1.21 at week 24. At week 27, the rabbits again exhibited an osteoporosis phenotype, reflected by a T-score of -2.55 ± 1.18 (**Figure 5D**). This result combined with all of the data provided in **Tables S3 - S5** demonstrates the reproducibility of density loss, partial recovery, and re-induction of osteoporosis which is achievable by repeated GC exposure providing a controlled framework for timing the VA procedure.

**Figure 5.**
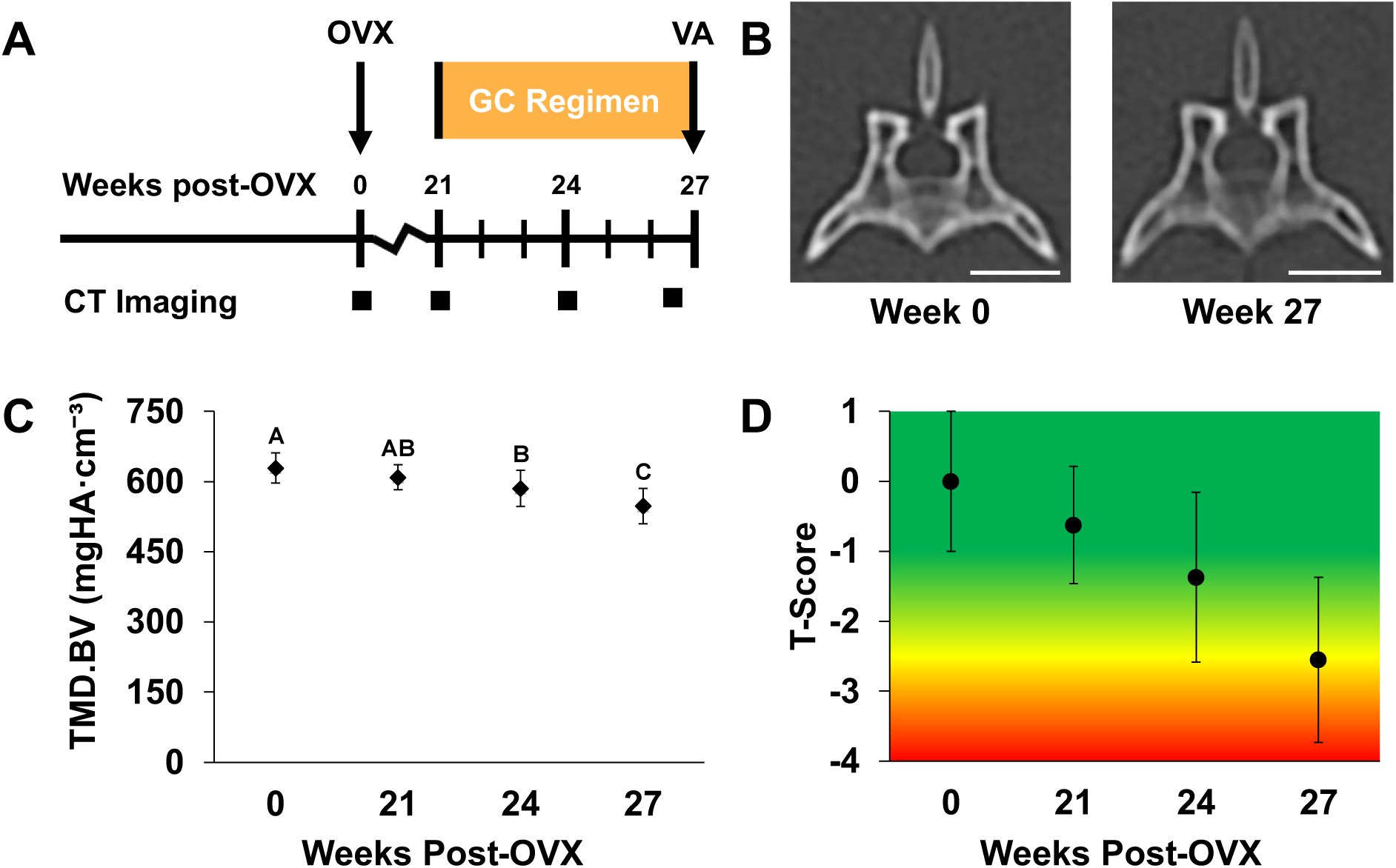
Osteoporosis Induction and Monitoring in Lumbar Vertebra. **(A)** Abridged study timeline showing experimental milestones. **(B)** Representative New Zealand White rabbit transverse L5 CT images pre-induction (Week 0) and post-induction (Week 27) show trabecular BMD reduction and cortical thinning. All scale bars indicate 2 cm. **(C)** Longitudinal analysis of TMD.BV (mgHA·cm⁻³) for pooled L4 - L6 vertebrae. **(D)** T-scores were derived from the mean and standard deviation of healthy (Week 0) TMD.BV (mgHA·cm⁻³) of pooled L4 - L6 measurements and used to elucidate healthy (-1 to 1), osteopenic (-1 to -2.5), and osteoporotic (< -2.5) bone phenotypes. Connecting letters indicate statistical groupings based on Tukey’s HSD post-hoc tests; values not sharing the same letter differ significantly (p < 0.05).

An open surgical VA approach consistently provided access to the lumbar vertebral bodies and intraoperative imaging verified alignment and safety throughout the procedure. As shown in **Figure 2A**, exposure of the L4 - L5 vertebral segment was obtained by a dorsal midline incision with careful retraction of paraspinal musculature, which provided reproducible visualization of the target anatomy. Creation of bilateral decorticated portals, illustrated in **Figure 2B**, enabled controlled entry through the dorsal cortex while avoiding pedicle disruption. Fluoroscopic verification ensured that trajectories directed toward the centrum were accurate and without cortical breach, as confirmed in **Figure 2C, Figure S3A,** and **Figure S3B**. Following fluoroscopy verification, PMMA was injected in a sequential manner on one side followed by the contralateral side as shown in **Figure 3A**. Final fluoroscopic images and video in **Figure 3B, Figure S3C,** and **Figure S3D** demonstrated bilateral PMMA distribution restricted to the cancellous compartment, supporting the reproducibility of augmentation while minimizing extraosseous spread. Earlier attempts to employ a percutaneous approach were unsuccessful due to limitations in trocar needle gauge and the anatomical constraints of the rabbit lumbar spine. Also, a single-side drilling approach frequently led to cement leakage into the spinal canal (**Figure S4A**), venous system, or other extraosseous regions (**Figure S4B**), most likely due to pressure buildup within the confined trabeculae. This was further limited by the small volume of injectable material that could be safely administered. The adoption of bilateral decorticated portals and reducing overall injection volume addressed these complications by offering dual access channels, which allowed for higher stable injection volumes while lowering intravertebral pressure and providing secondary space for material overflow. This adaptation mirrors clinical strategies in percutaneous vertebroplasty (PVP) and kyphoplasty (PKP), where bilateral access is often favored to optimize cement distribution and limit the risk of leakage. [44, 45] Collectively, the surgical procedure presented in **Figure 2** and **Figure 3** confirms that bilateral VA can be performed safely and reproducibly in an osteoporotic rabbit lumbar spine model. The integration of open surgical access with fluoroscopic verification of portal placement establishes a workflow that parallels clinical VA practice and supports this model as a relevant platform for preclinical evaluation of injectable biomaterials in osteoporotic vertebrae.

Following establishment of this clinically relevant workflow, VA outcomes were assessed both qualitatively and quantitatively. Evaluation of injected PMMA volume with qCT excitingly showed similar PMMA injection volumes (**Figure 6A**). Specifically, an average of 37.47 ± 9.44 mm³ and 31.11 ± 10.28 mm³ was injected into the L4 and L5, respectively, based on the five rabbits that survived the procedure. Notably, when the injected PMMA volume exceeded approximately 100 µL (*), hind limb paralysis was observed, thus the effective upper limit for safe injections was found to be around 50 µL (**Figure 6A, Table S6**). L6 showed no observable BaSO_4_ contrast agent corresponding to a lack of PMMA as expected. When comparing post-VA qCT parameters holistically across L4 - L6 (**Table S6**), no statistically significant differences were observed between groups for any of the measured whole, trabecular, and cortical qCT bone parameters. This suggests that, when PMMA volume is excised from analysis, the overall bone quality and structure remained consistent across injected (L4 and L5) and non-injected vertebrae (L6), indicating a minimal impact of VA on adjacent qCT-assessed bone properties. When pooling the injected vertebrae (L4 and L5) to evaluate pre-VA versus post-VA differences (**Table S7**), most trabecular and cortical parameters remained stable, further underscoring minimal overall impact of VA on overall bone microarchitecture. For instance, TMD.BV increased only slightly from 537.41 ± 35.68 mgHA·cm⁻³ to 553.70 ± 36.59 mgHA·cm⁻³, while BV/TV rose modestly from 0.895 ± 0.023 to 0.909 ± 0.015, with both changes being statistically insignificant. Similarly, cortical indices such as Ct.Ar (61.36 ± 6.89 mm² to 63.83 ± 6.56 mm²), Ct.Th (1.09 ± 0.25 mm to 1.24 ± 0.21 mm), and Ct.Po (0.28 ± 0.09 to 0.24 ± 0.06) remained unaffected, confirming that the developed VA procedure exerts minimal measurable effects on the cortical and trabecular compartments. In contrast, bone surface (BS) parameters were significantly reduced for injected vertebrae from 1609 ± 320 mm² to 1365 ± 232 mm² (**Figure 6B** and **Table S7**). Parameters derived from BS such as BS/TV declined from 0.660 ± 0.162 mm⁻¹ to 0.555 ± 0.123 mm⁻¹ and BS/BV fell from 0.740 ± 0.191 mm⁻¹ to 0.611 ± 0.136 mm⁻¹, both significantly reduced. This BS decline was not observed in L6 vertebra, which had a more stable BS of 1505 ± 270 and 1388 ± 271 mm² (**Figure 6C** and **Table S8**) as well as derived BV/TV values of 0.912 ± 0.017 and 0.923 ± 0.013 for pre-VA and post-VA, respectively (**Table S8**). This selective reduction in surface-related metrics can be directly attributed to the mildly disruptive VA surgical procedure. During augmentation, drilling burs are used to create a cortical portal to access trabecular bone. The subsequent injection of PMMA cement into this prepared trabecular space further fills and smooths the local architecture, displacing bone surfaces and lowering the surface-to-volume ratio due to exclusion of PMMA in qCT analysis. This observation can be seen in **Figure 6D**, which shows PMMA highlighted in a blue overlay within three-dimensional lateral reconstruction, as well as in the corresponding two-dimensional coronal, sagittal, and transverse plane. These results reinforce that VA does not appreciably alter trabecular thickness, bone volume fraction, connectivity density, or cortical indices, but has a clear and expected impact on whole bone surface metrics.

**Figure 6.**
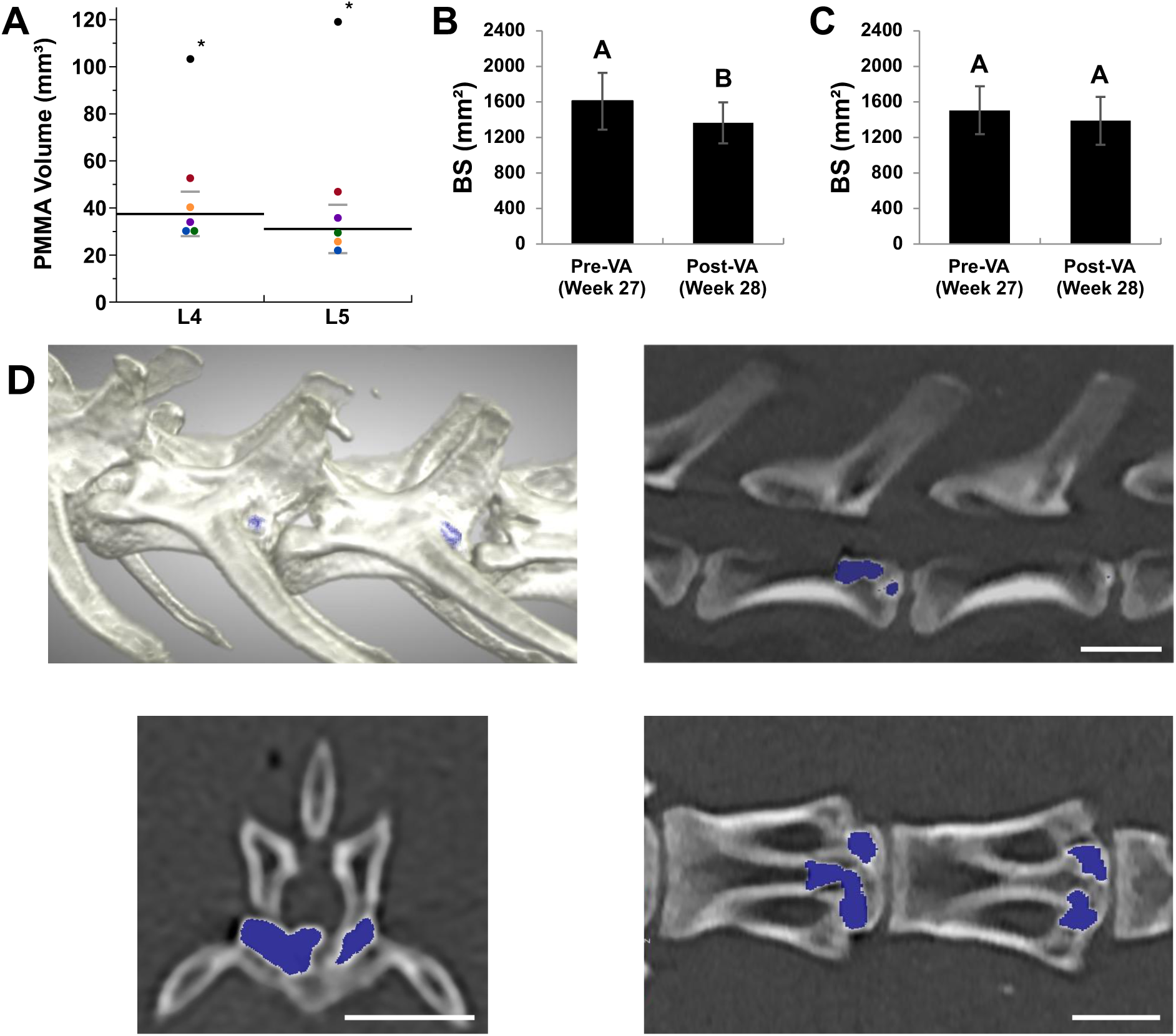
Summary of Vertebral Augmentation (VA) Outcomes. **(A)** Quantitative assessment of injected cement volume in treated vertebrae (L4 and L5) using qCT analysis. Mean injected PMMA volumes were 37.47 ± 9.44 mm³ (L4) and 31.11 ± 10.28 mm³ (L5), indicating comparable injection volumes across levels with each rabbit represented by a different color. Data points indicated with (*) denotes a rabbit that survived VA but later developed hind limb paralysis and was euthanized, so it was not included in later analysis. **(B)** Bone surface (BS) measurements for injected vertebrae (L4 and L5) before (Week 27) and after VA (Week 28). **(C)** Bone surface (BS) measurements for a non-injected vertebra (L6) before (Week 27) and after VA (Week 28). **(D)** Representative post-VA qCT images of injected L4 - L5 vertebrae, including three-dimensional lateral reconstructions and two-dimensional coronal, sagittal, and transverse planes. PMMA (blue overlay) is localized within the trabecular compartment, illustrating cement distribution following surgical access and material injection. All scale bars indicate 2 cm. Values are presented as mean ± standard deviation. Groups that possess different letters have statistically significant difference in mean (p ≤ 0.05) whereas those that possess the same letter have similar means (p > 0.05).

### Conclusion

In this study, we successfully developed and validated an osteoporotic rabbit model of vertebral augmentation that combines reproducible osteoporosis induction with a clinically inspired, fluoroscopically guided surgical workflow. The OVX + glucocorticoid regimen consistently produced quantifiable osteoporotic bone loss, as confirmed by qCT-derived reductions in mineral density, T-scores ≤ -2.5, and controlled microarchitectural deterioration. The open dorsal bilateral-access VA procedure allowed for the safe and reproducible delivery of PMMA into the L4 and L5 lumbar vertebrae with minimal cement leakage which was readily confirmed by fluoroscopy. Quantitative CT analysis showed that while VA predictably reduced certain bone surface-related parameters due to cortical portal creation and cement filling, all other trabecular and cortical indices remained stable, indicating that bone tissue was structurally preserved overall.

By integrating disease-relevant bone degeneration with a clinically aligned vertebral augmentation technique, this model bridges the translational gap between small rodent and large-animal systems. It offers an accessible, cost-effective, and translationally relevant platform to test emerging biomaterials for spinal regeneration applications. Ultimately, the establishment of this model lays the groundwork for the development of alternative materials to PMMA that may improve both safety and regenerative outcomes for patients suffering from osteoporotic vertebral compression fractures.

## Author Contributions (CRediT)

Conceptualization and methodology were contributed by AH, RS, DM, JHL, and BU; investigation and data curation were performed by AH, AG, AK, JW, SH, SR, and JHL; writing—original draft was completed by AH, JHL, and BU; writing—review and editing were carried out by all authors; resources were provided by RS, DM, and JHL; funding acquisition was led by BU; and supervision was provided by RS, DM, and BU.

