## Supplementary Information for "Filling the Spinal Fracture Treatment Gap: An Osteoporotic Rabbit Model of Vertebral Augmentation"

**Supplementary Materials for Filling the Spinal Fracture Treatment Gap: An Osteoporotic Rabbit Model of Vertebral Augmentation**

August J. Hemmerla^1^, Abigail R. Grisolano^1^, Austin D. Kimes^1,2^, John T. Wray^2^, Sam E. Huddleston^1^, Shwetha Ramachandra^1^, Ryan E. Schultz^3^, Don K. Moore^4^, Ji-Hey Lim^5^, and Bret D. Ulery^1,6,7,#^

^1^Department of Chemical and Biomedical Engineering, University of Missouri, Columbia, MO 65211, USA

^2^College of Veterinary Medicine, University of Missouri, Columbia, MO 65211, USA

^3^Department of Radiology, University of Missouri, Columbia, MO 65212, USA

^4^Department of Orthopedic Surgery, University of Missouri, Columbia, MO 65201, USA

^5^Department of Veterinary Neurology and Neurosurgery, University of California, Davis, Davis, CA 95616, USA

^6^NextGen Precision Health Institute, University of Missouri, Columbia, MO 65211, USA

^7^Materials Science and Engineering Institute, Columbia, MO 65211, USA

| 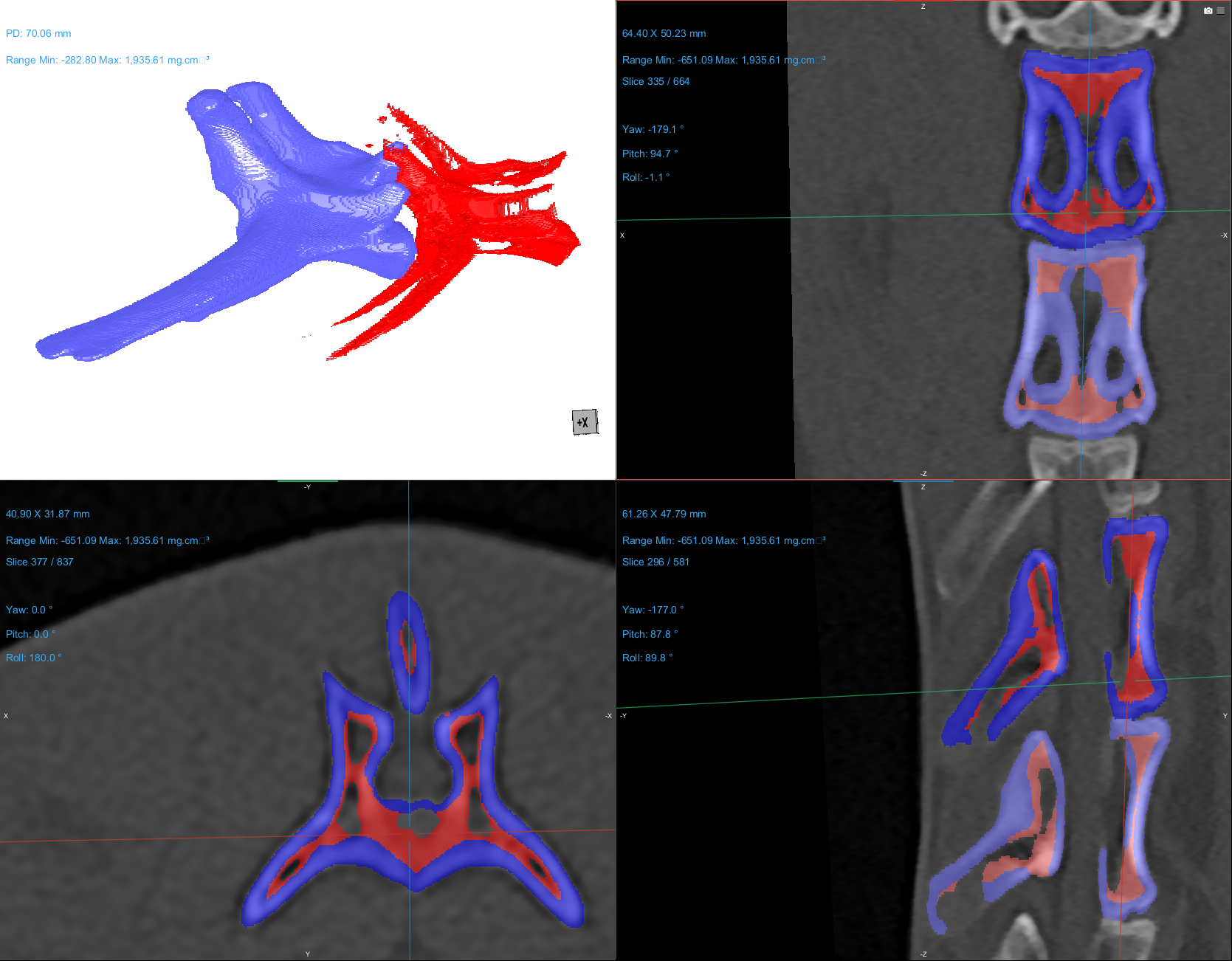  **A**  **B** | | 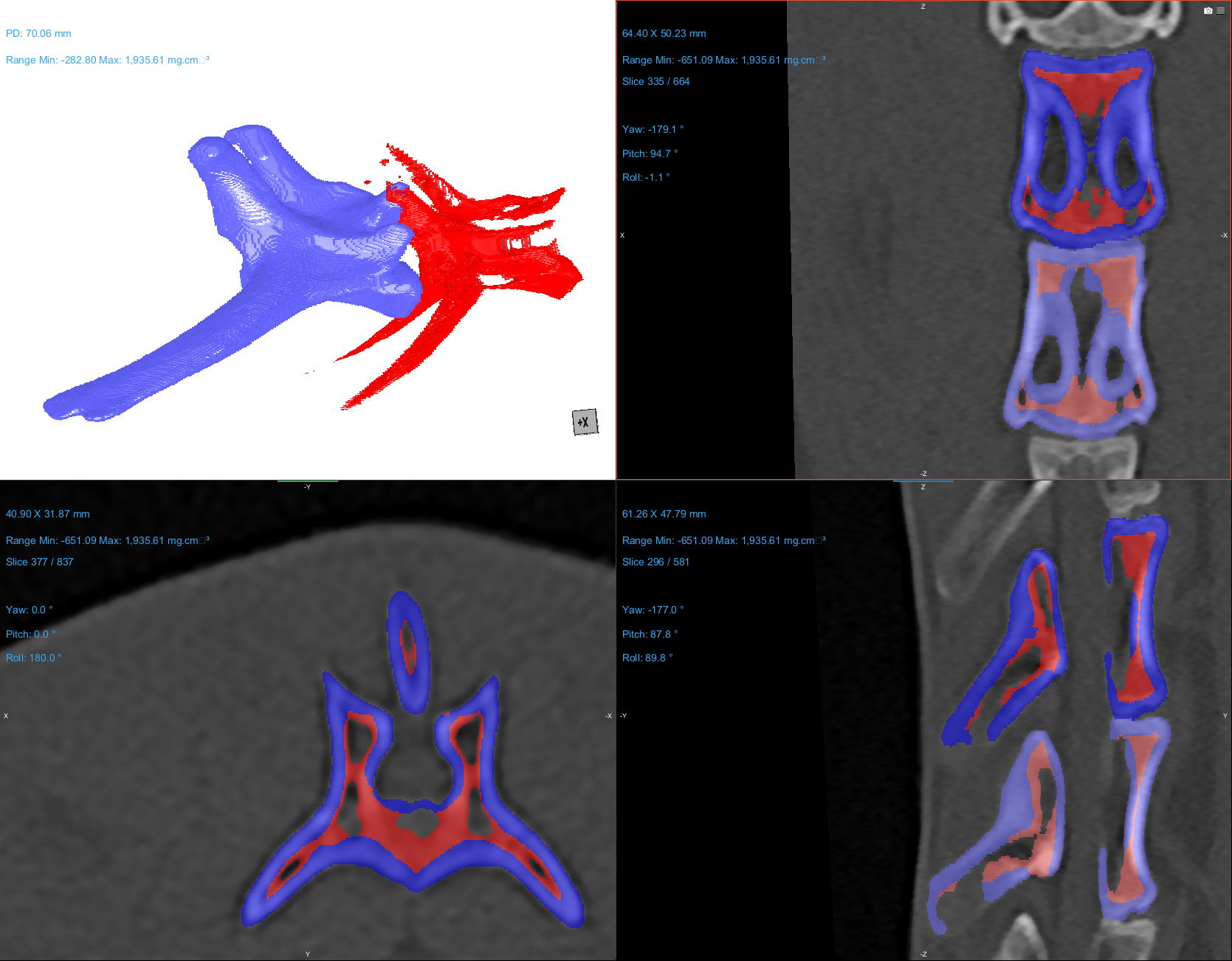 |
| --- | --- | --- |
| 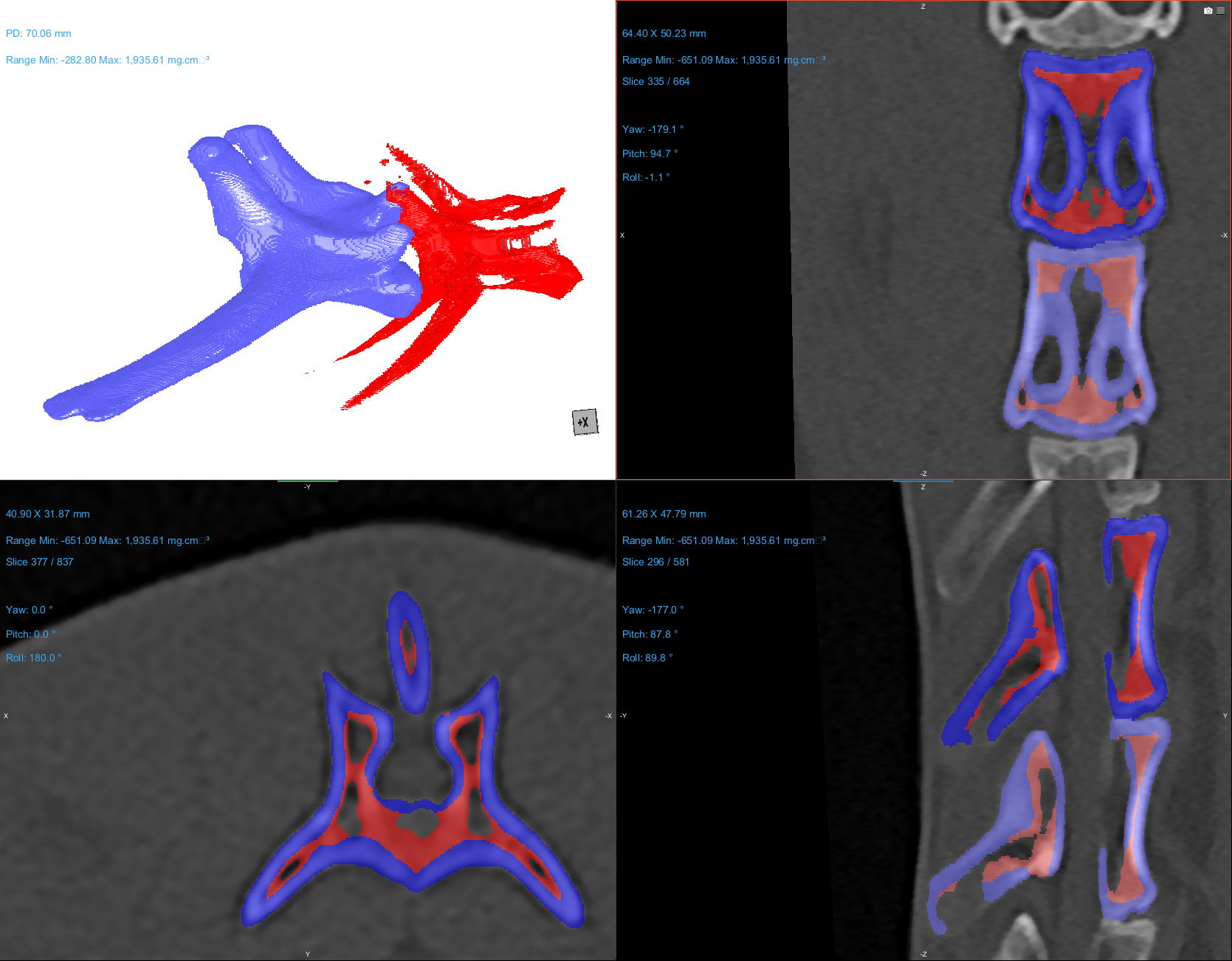  **C** | 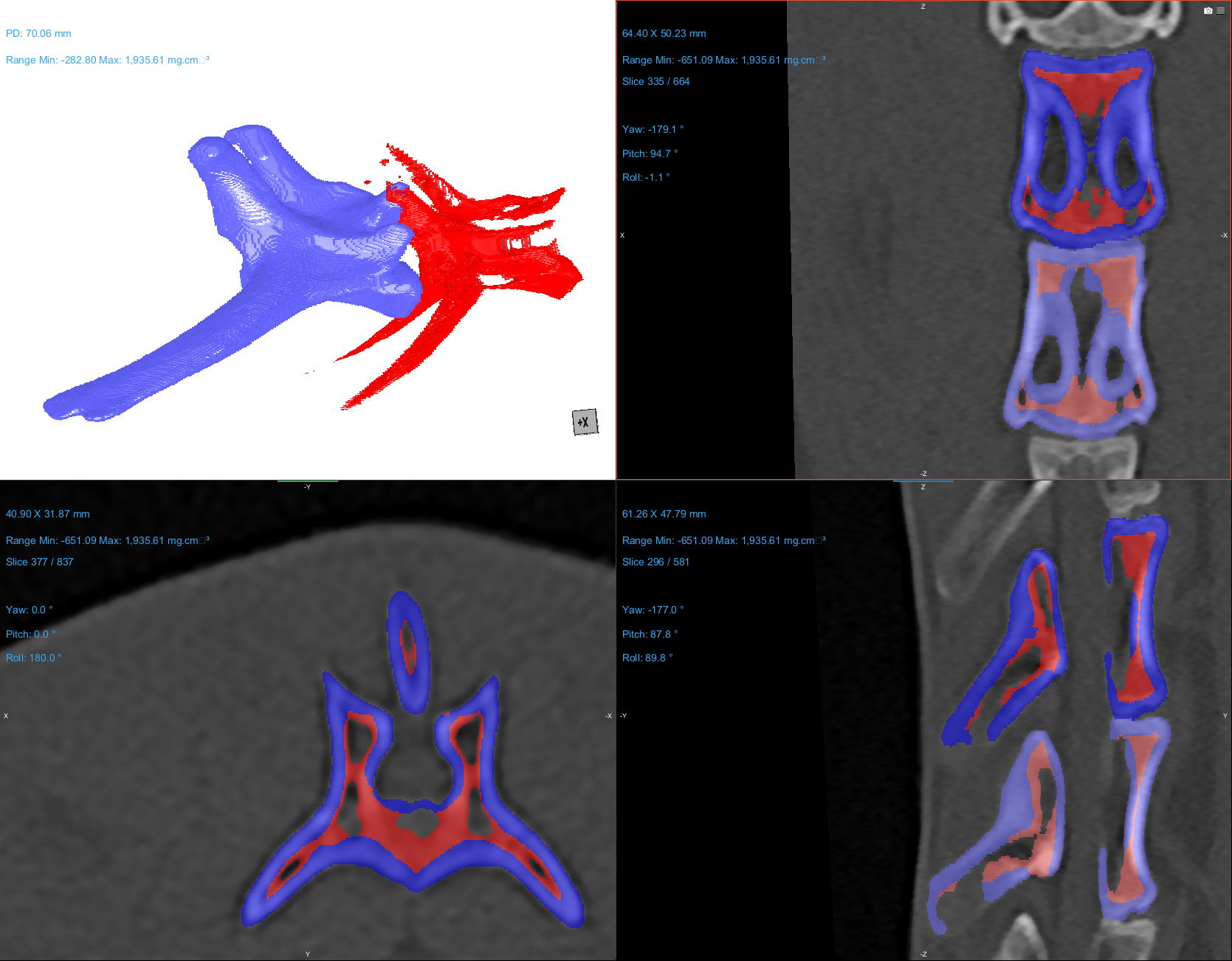  **D** | |
| **Figure S1.** **Representative quantitative computed tomography (qCT)-based segmentation of the lumbar vertebrae in a New Zealand White rabbit.** **(A)** A three-dimensional lateral reconstruction of L4 - L5 demonstrates cortical (blue) and trabecular (red) bone compartment segmentation. With additional views demonstrating the **(B)** transverse, **(C)** sagittal, and **(D)** coronal planes. | | |

|  | **Pre-OVX (Week 0)** | **Pre-VA (Week 27)** |
| --- | --- | --- |
| **L4 Transverse** | 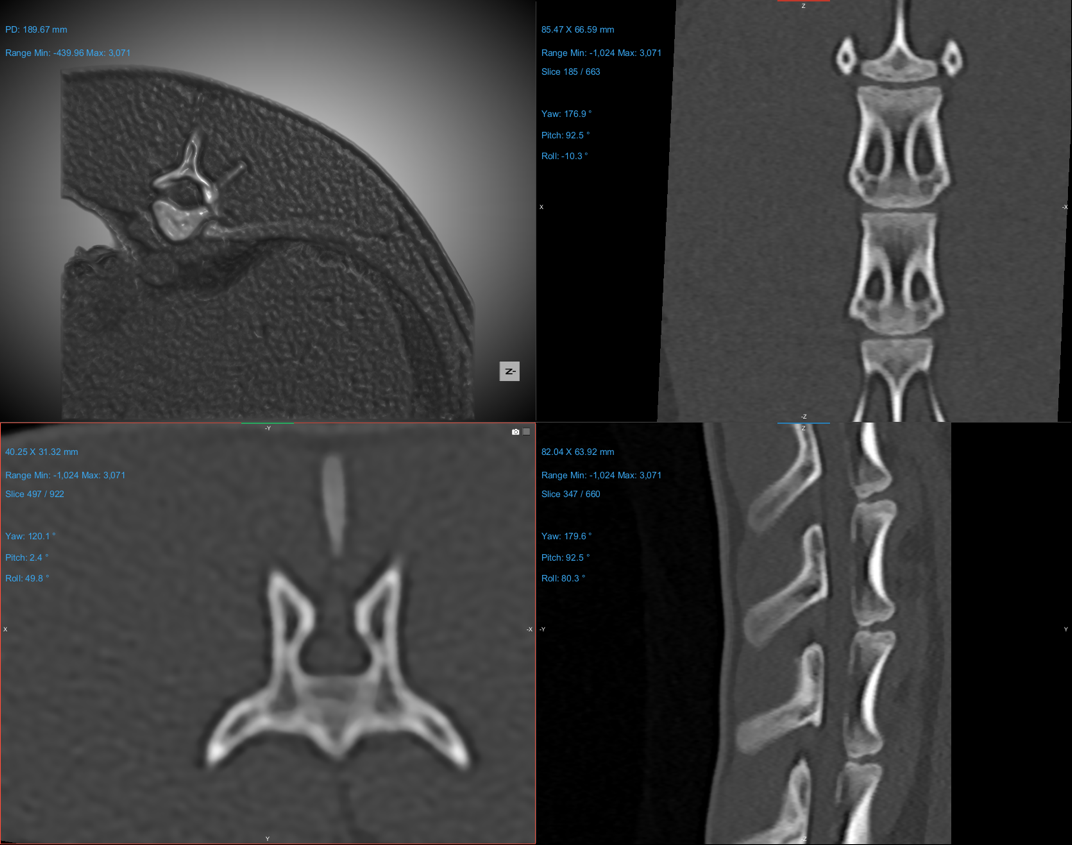 | 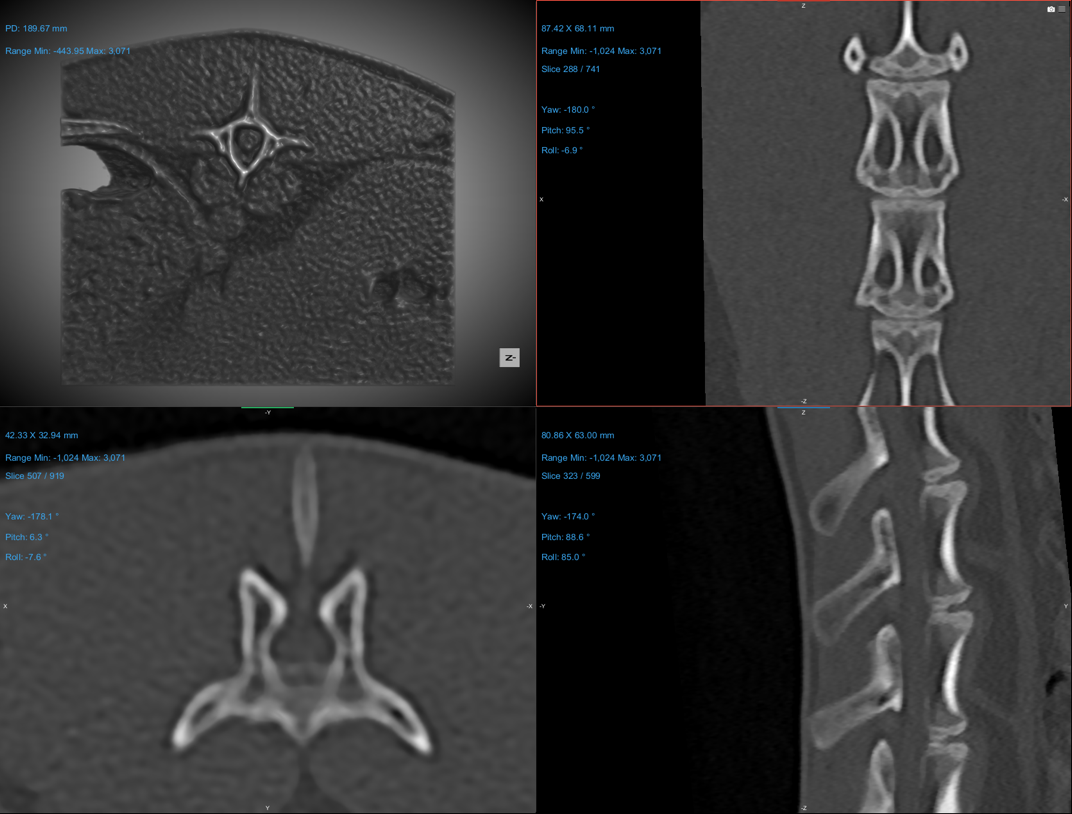 |
| **L5 Transverse** | 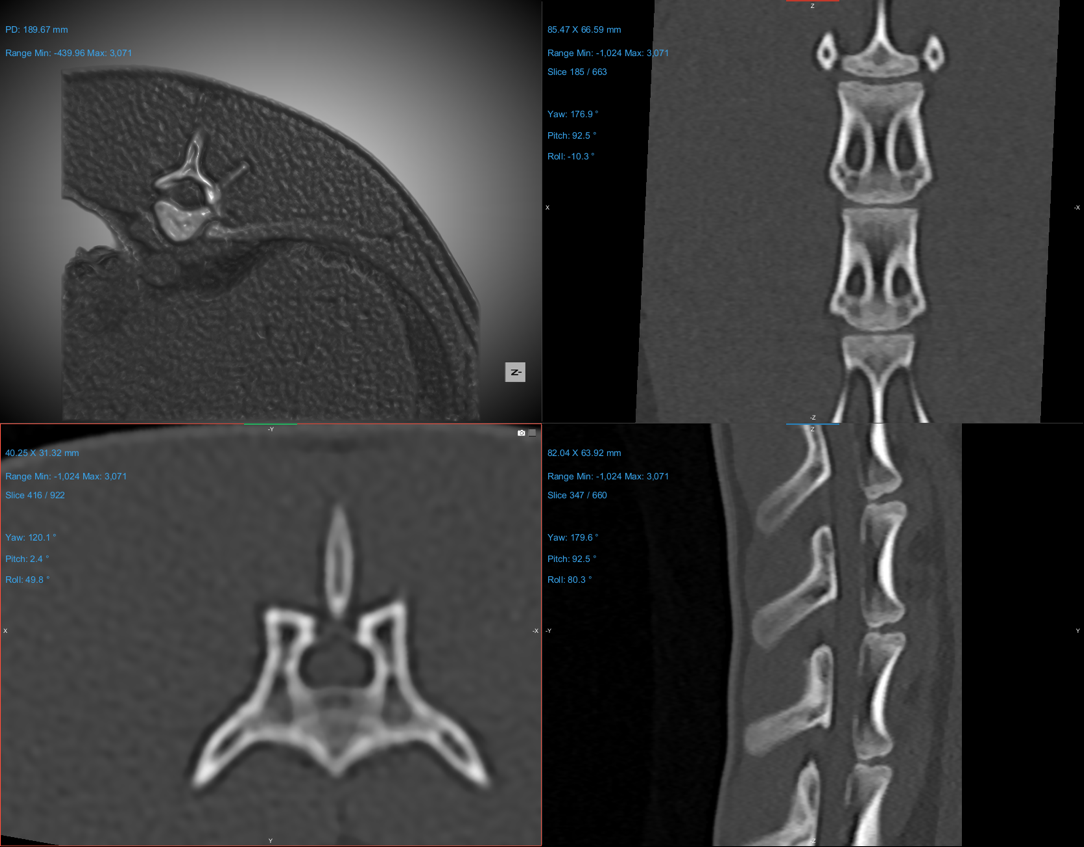 | 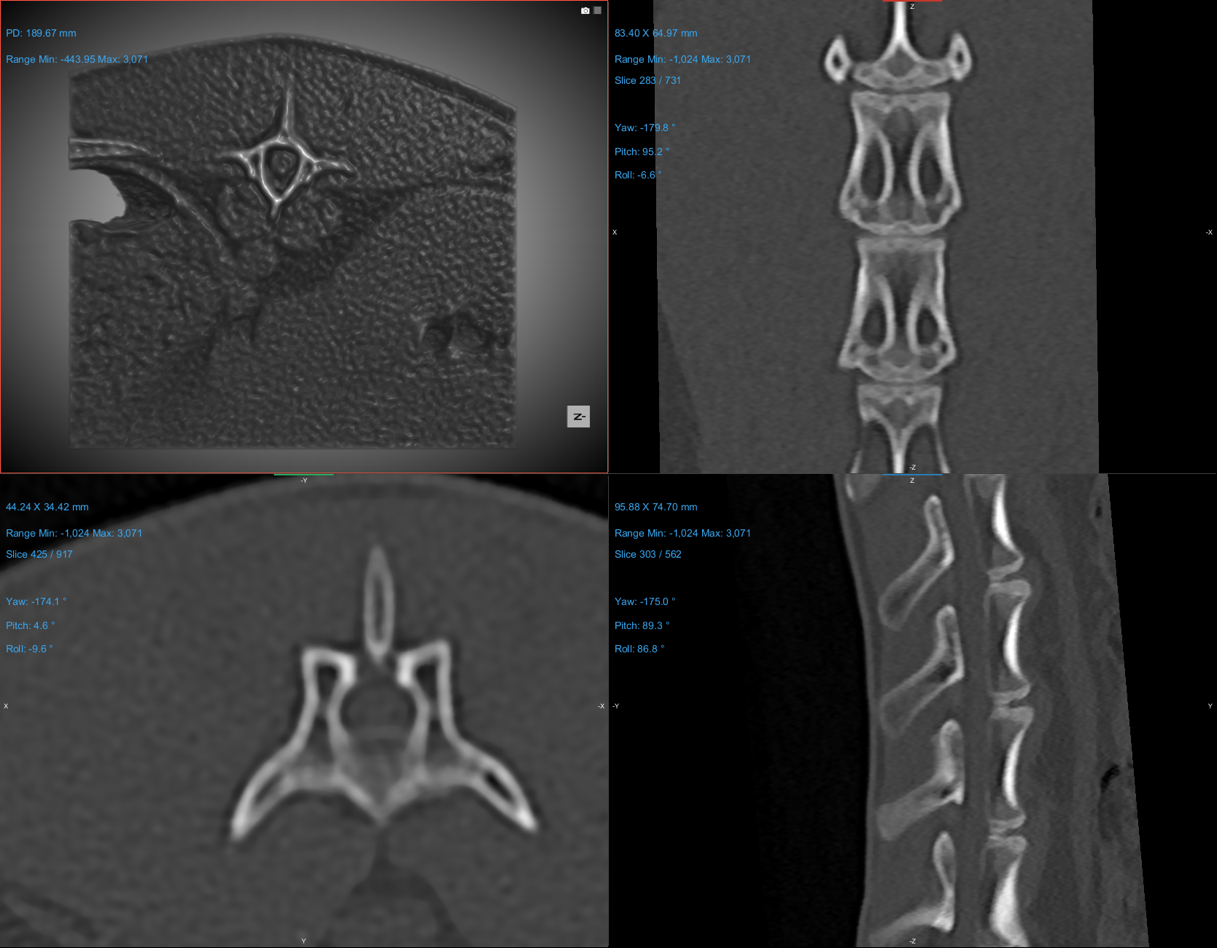 |
| **L4 – L5 Sagittal** | 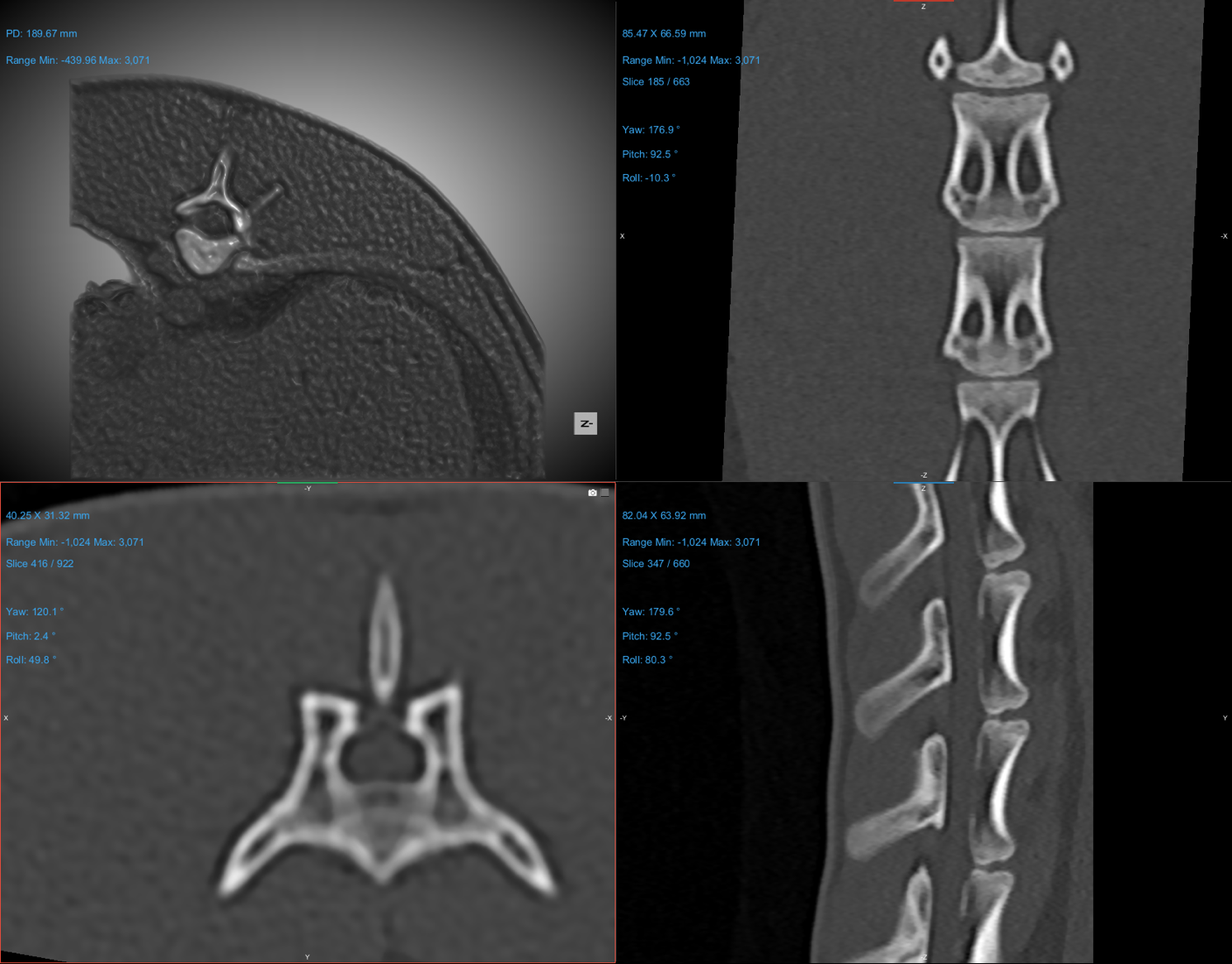 | 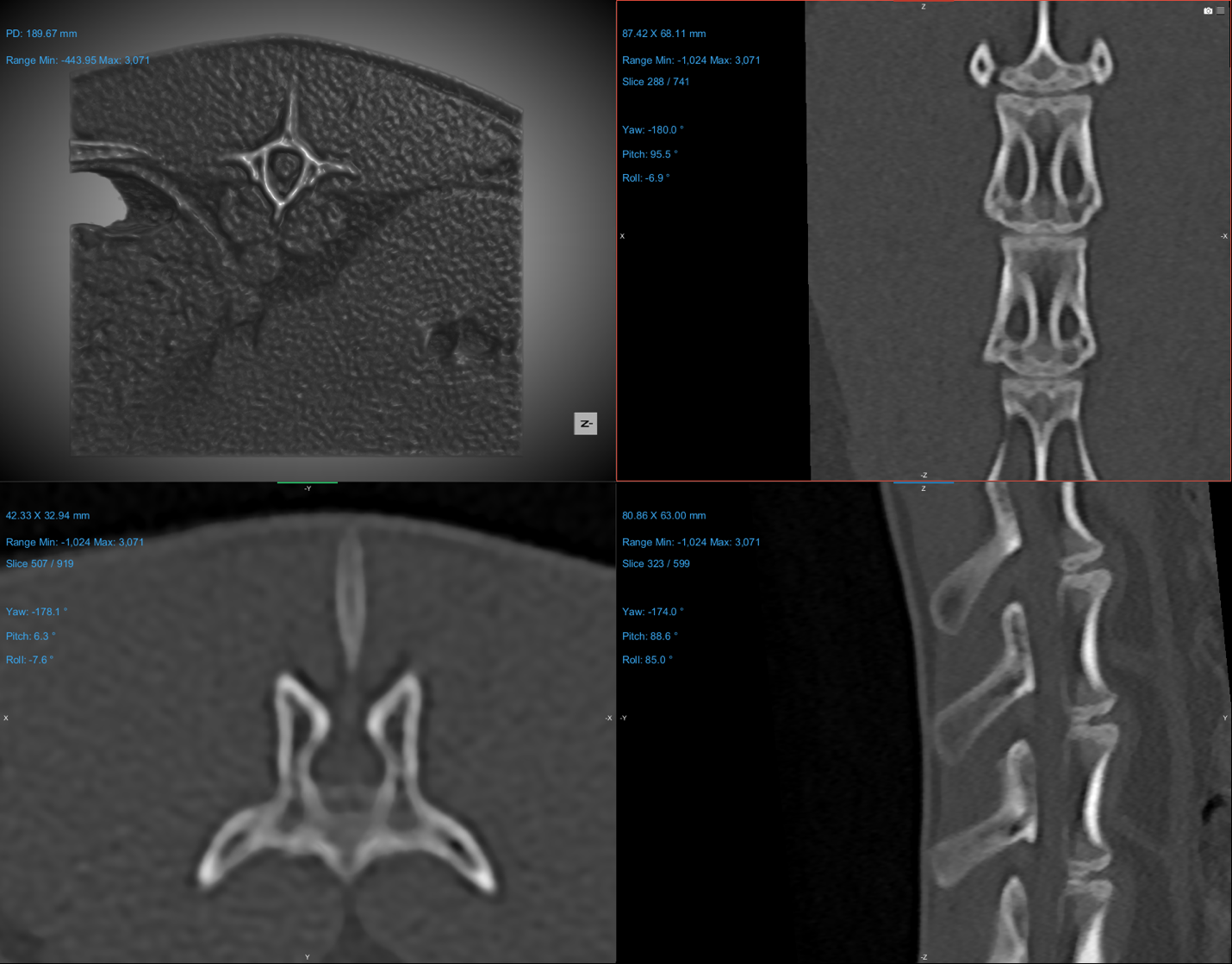 |
| **L4 – L5 Coronal** | 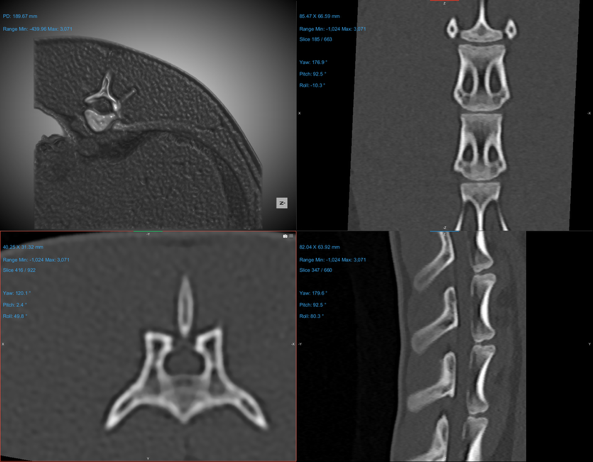 | 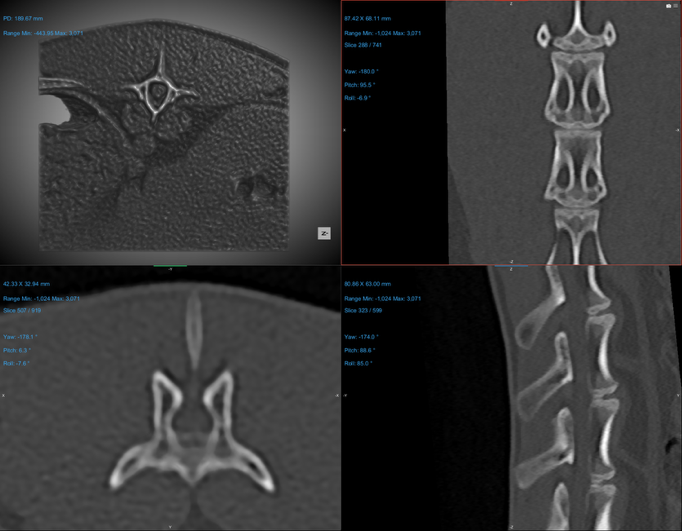 |
| **Figure S2.** **Representative CT Images of Lumbar Vertebrae in New Zealand White Rabbits Demonstrating Osteoporosis Progression at Week 0 and Week 27. (A)** L4 transverse, **(B)** L5 transverse, **(C)** sagittal, and **(D)** coronal planes. Progressive reductions in trabecular bone density and cortical thinning are evident over time, indicating structural deterioration associated with osteoporosis development. | | |

| **A** | **Lateral View**  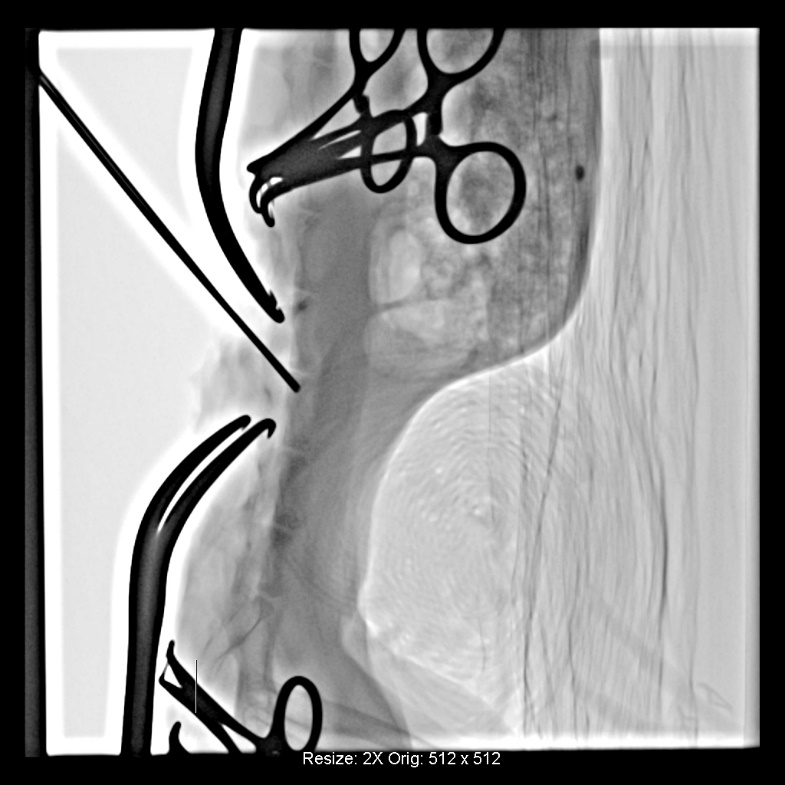 | **Dorsoventral (DV) View** 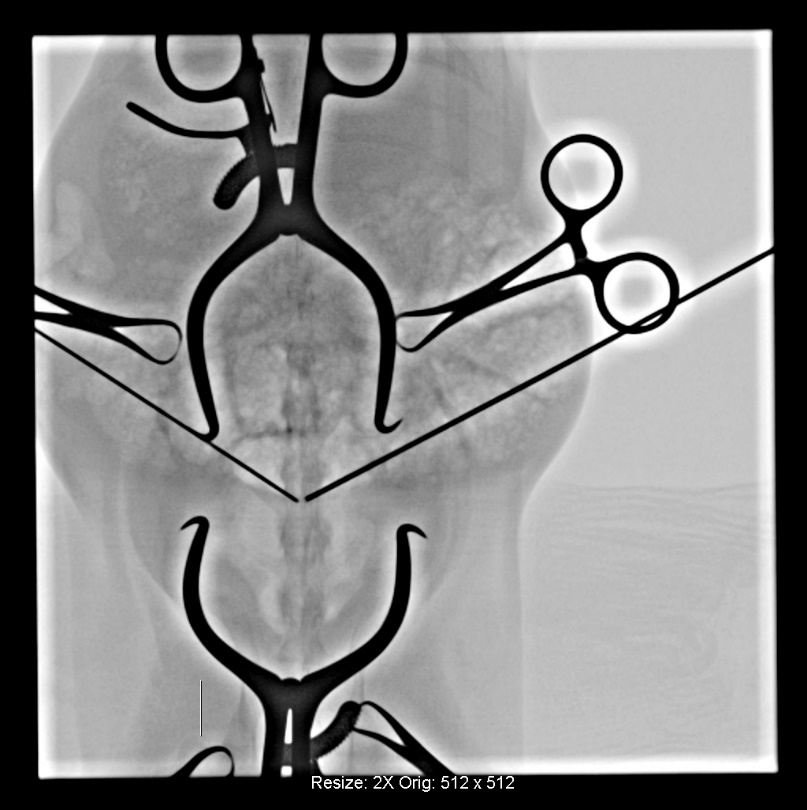 |
| --- | --- | --- |
| **B** | 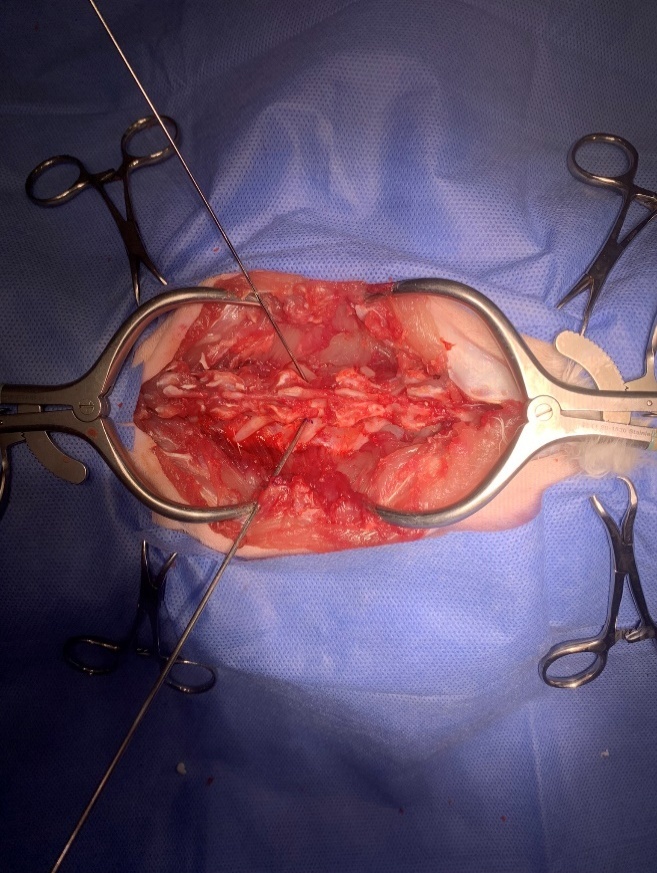 | |
| **C** | **Lateral View**  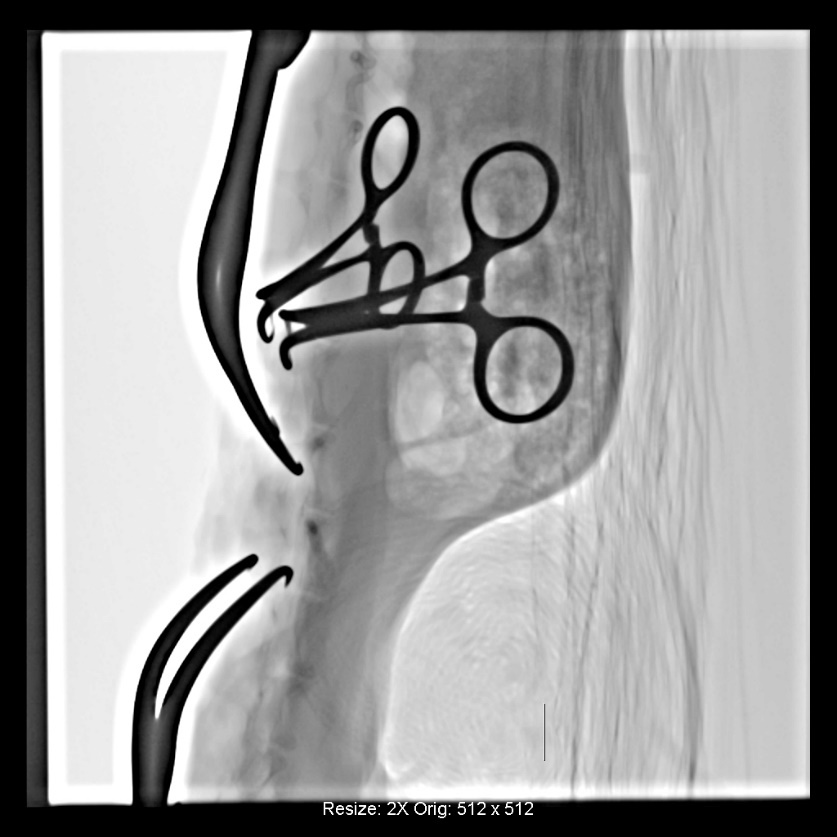 | **Dorsoventral (DV) View**  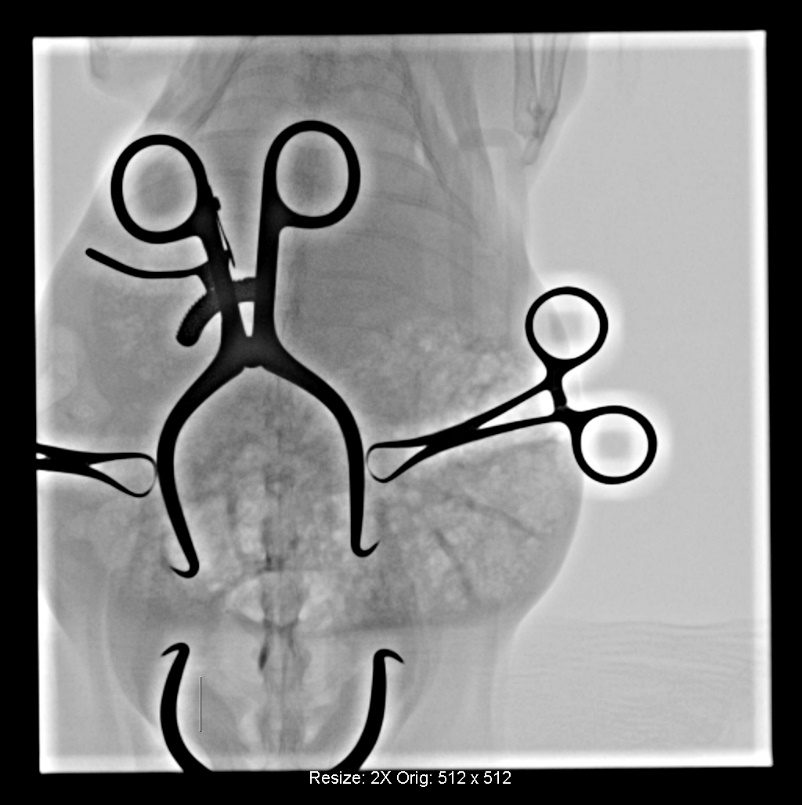 |
| **D** | 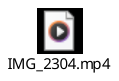 | |
| **Figure S3. Example of bilateral injection demonstrating the procedural sequence and outcomes. (A)** Lateral and dorsoventral post-portal drilling using fluoroscopic imaging allows for the verification of injection depth. **(B)** A corresponding intraoperative surgical image showing the same dorsoventral plane as the second imagine in Part A. **(C**) Lateral and dorsoventral post-injection fluoroscopic images confirm symmetric bilateral PMMA distribution **(D)** A post-injection video illustrates the two established portals and adequate cement delivery. | | |

|  | **Lateral View** | **Dorsoventral (DV) View** |
| --- | --- | --- |
| **A** | 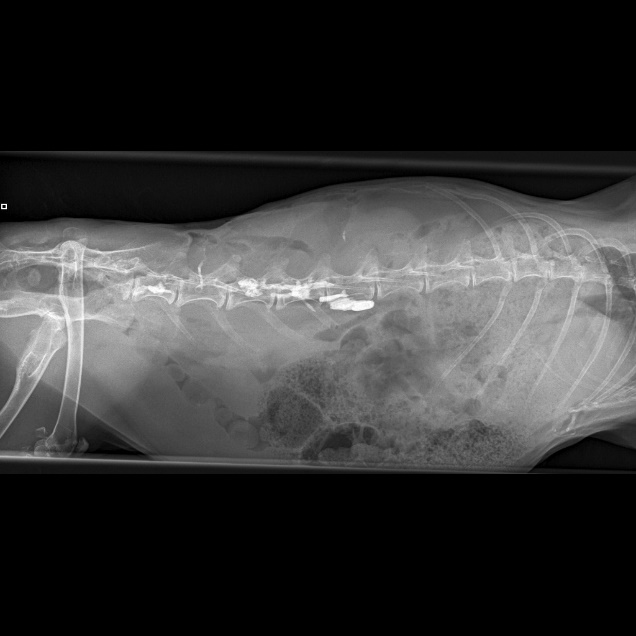 | 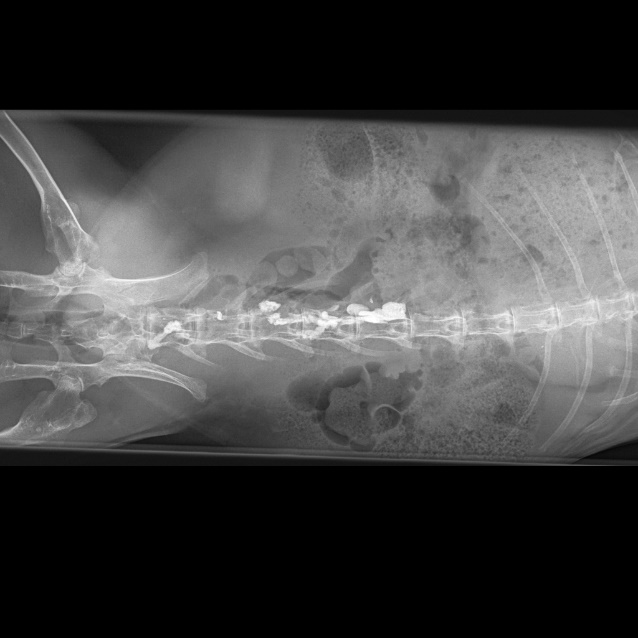 |
| **B** | 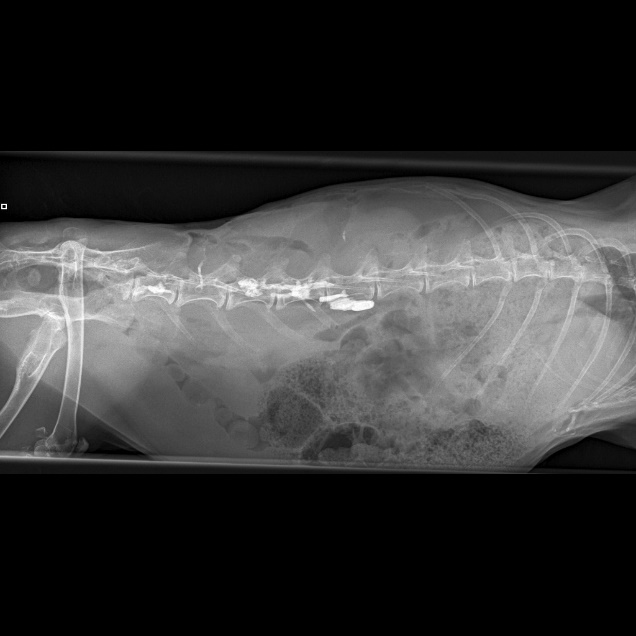 | 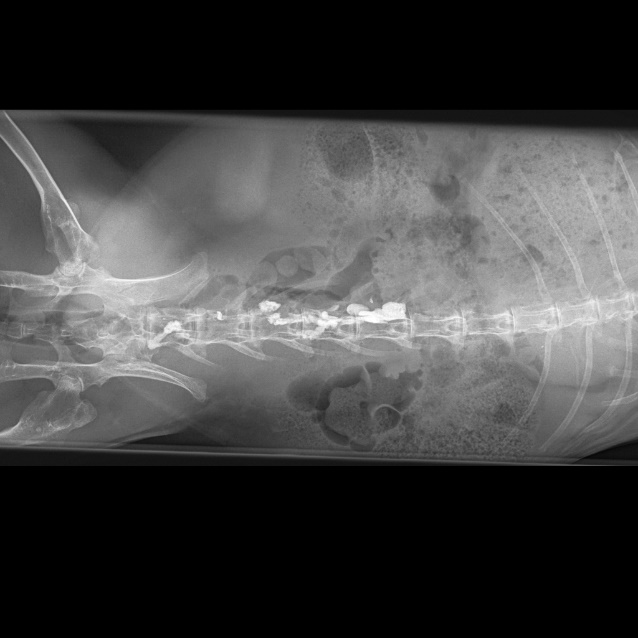 |
| **Figure S4. Example fluoroscopic images demonstrating complications associated with single-side drilling. (A)** Cement leaked into the spinal canal and **(B)** extraosseous leakage extended well beyond the vertebral boundary. | | |

| **Table S1. Variables, Units, and Descriptions of Whole-Bone, Trabecular, and Cortical qCT Parameters.** | | | |
| --- | --- | --- | --- |
|  | **qCT Parameters** | **Units** | **Description** |
| **Whole Bone** | Tissue Mineral Density (Bone Volume) (**TMD.BV**) | mgHA·cm⁻³ | Mean tissue mineral density of bone tissue across cortical and trabecular region of interests (ROIs); material density of bone itself (excluding soft tissue) |
|  | Total Volume (**TV**) | mm³ | Volume of the filled bone ROI |
|  | Bone Volume (**BV**) | mm³ | Volume of segmented bone (cortical and trabecular) within the ROI |
|  | Bone Surface (**BS**) | mm² | Surface area of the segmented bone ROI computed via marching cubes or weighted-voxel approximation |
|  | Bone Volume Fraction (**BV/TV**) | — | Ratio of bone volume to total volume within the ROI |
|  | Bone Surface Density (**BS/TV**) | mm⁻¹ | Ratio of bone surface area to total volume (surface density) |
|  | Specific Bone Surface (**BS/BV**) | mm⁻¹ | Ratio of bone surface area to bone volume (specific surface) |
| **Trabecular** | Bone Mineral Density (**BMD.Me**) | mgHA·cm⁻³ | Mean bone mineral density of the mixed region comprised of trabecular bone and the medullary cavity (mixed bone–soft tissue region) |
|  | Tissue Mineral Density (Trabecular Bone) (**TMD.Tb**) | mgHA·cm⁻³ | Mean tissue mineral density measured within the trabecular bone ROI; material density of trabecular bone (excluding soft tissue) |
|  | Average Trabecular Thickness (**Tb.Th**) | mm | Average local thickness of trabeculae volume based on the diameter of the largest inscribed sphere at each point (from thickness map) |
|  | Connectivity Density (**Conn.D**) | mm⁻³ | Connectivity of the trabecular network was calculated as (1 - Euler–Poincaré Characteristic) normalized to trabecular bone volume, providing the number of connected trabecular elements per unit volume. [1] |
|  | Anisotropy (SVD) | — | Degree of orientation of trabecular architecture using the Star Volume Distribution method [2] |
| **Cortical** | Tissue Mineral Density (Cortical bone) (TMD.Co) | mgHA·cm⁻³ | Mean tissue mineral density within the cortical bone ROI; material density of cortical bone (excludes soft tissue) |
|  | Average Cortical Area (**Ct.Ar**) | mm² | Average cross-sectional area of cortical bone within the selected bone length |
|  | Average Cortical Thickness (**Ct.Th**) | mm | Average local thickness of cortical bone defined by maximal inscribed sphere diameter at each point (from thickness map) [3] |
|  | Average Total Area (**Tt.Ar**) | mm² | Cross‑sectional area of filled bone (cortical and marrow) computed as total volume divided by bone length |
|  | Cortical Porosity (**Ct.Po**) | — | Ratio of pore volume to cortical bone volume within the cortical ROI |
|  | Periosteal Perimeter (**Ps.Pm**) | mm | Average perimeter length of the periosteal boundary within the selected bone length |
|  | Periosteal Surface (3D) (**Ps.S3D**) | mm² | Total surface area of the periosteum computed with a weighted‑voxel approximation |

| **Table S2. Healthy Lumbar Comparison of qCT Parameters.** An overview of general qCT parameters for L3 - L6 with values is presented as mean ± standard deviation. Across a parameter (*i.e.*, row), groups that possess different letters have statistically significant difference in mean (p ≤ 0.05) whereas those that possess the same letter have similar means (p > 0.05). N = 8 (total vertebrae assessed per group). | | | | |
| --- | --- | --- | --- | --- |
| **Week 0** | **L3** | **L4** | **L5** | **L6** |
| **TMD.BV (mgHA·cm⁻³)** | 610.7 ± 30.3 (A) | 614.6 ± 31.8 (A) | 625.2 ± 25.4 (A) | 647.5 ± 32.3 (A) |
| **TV (mm³)** | 2043 ± 171(A) | 2240 ± 116 (AB) | 2406 ± 119 (B) | 2387 ± 184 (B) |
| **BV (mm³)** | 1877 ± 153 (A) | 2072 ± 102 (B) | 2236 ± 105 (B) | 2245 ± 166 (B) |
| **BV/TV** | 0.919 ± 0.008 (A) | 0.925 ± 0.010 (A) | 0.929 ± 0.008 (AB) | 0.940 ± 0.012 (B) |
| **TMB.Tb (mgHA·cm⁻³)** | 536.0 ± 76.8 (A) | 501.4 ± 46.1 (A) | 492.6 ± 51.2 (A) | 512.2 ± 63.0 (A) |
| **Tb.Th (mm)** | 1.552 ± 0.200 (A) | 1.558 ± 0.100 (A) | 1.550 ± 0.103 (A) | 1.502 ± 0.120 (A) |
| **TMD.Co (mgHA·cm⁻³)** | 616.5 ± 57.1 (A) | 642.0 ± 41.8 (A) | 653.3 ± 30.5 (A) | 674.3 ± 37.3 (A) |
| **Ct.Th (mm)** | 1.164 ± 0.348 (A) | 1.245 ± 0.245 (A) | 1.313 ± 0.331 (A) | 1.326 ± 0.270 (A) |

| **Table S3. Comparison of Pooled Lumbar Vertebrae (L4 - L6) qCT Parameters Between Week 0 and Week 8.** During the period following OVX and the initial glucocorticoid (GC) regimen, animals were able to reach an osteoporotic state. Bone mineral density measurements (*i.e.*, TMD.BV, TMD.Tb, TMD.Co, and BMD.Me) declined considerably by Week 8, while geometry (*i.e.*, TV, BV, BS, Ct.Ar, Tt.Ar, and Ps.Pm) and most microarchitecture measurements (*i.e.*, Tb.Th, Conn.D, Anisotropy, and Ct.Po) remained stable. Values are presented as mean ± standard deviation where, across a parameter (*i.e.*, row), groups that possess different letters have statistically significant difference in mean (p ≤ 0.05) whereas those that possess the same letter have similar means (p > 0.05). N = 24 (total vertebrae assessed per group). | | | |
| --- | --- | --- | --- |
|  |  | **Week 0** | **Week 8** |
|  | **TMD.BV (mgHA·cm⁻³)** | 629.1 ± 31.9 (A) | 479.2 ± 38.5 (B) |
|  | **TV (mm³)** | 2344 ± 156 (A) | 2411 ± 228 (A) |
|  | **BV (mm³)** | 2184 ± 147 (A) | 2203 ± 224 (A) |
|  | **BS (mm²)** | 1192 ± 326 (A) | 1351 ± 743 (A) |
|  | **BV/TV** | 0.932 ± 0.012 (A) | 0.914 ± 0.041 (A) |
|  | **BS/TV (mm⁻¹)** | 0.511 ± 0.143 (A) | 0.567 ± 0.330 (A) |
|  | **BS/BV (mm⁻¹)** | 0.550 ± 0.157 (A) | 0.636 ± 0.390 (A) |
|  | **BMD.Me (mgHA·cm⁻³)** | 357.5 ± 79.9 (A) | 262.4 ± 85.9 (B) |
|  | **TMD.Tb (mgHA·cm⁻³)** | 502.0 ± 52.1 (A) | 380.6 ± 78.7 (B) |
|  | **Tb.Th (mm)** | 1.537 ± 0.106 (A) | 1.573 ± 0.266 (A) |
|  | **Conn.D (mm⁻³)** | 0.168 ± 0.137 (A) | 0.147 ± 0.176 (A) |
|  | **Anisotropy (SVD)** | 0.679 ± 0.122 (A) | 0.690 ± 0.186 (A) |
|  | **TMD.Co (mgHA·cm⁻³)** | 656.5 ± 37.7 (A) | 484.1 ± 56.1 (B) |
|  | **Ct.Ar (mm²)** | 65.16 ± 6.63 (A) | 66.98 ± 8.81 (A) |
|  | **Ct.Th (mm)** | 1.295 ± 0.274 (A) | 1.289 ± 0.533 (A) |
|  | **Tt.Ar (mm²)** | 65.17 ± 6.64 (A) | 66.98 ± 8.82 (A) |
|  | **Ct.Po** | 0.218 ± 0.068 (A) | 0.272 ± 0.176 (A) |
|  | **Ps.Pm (mm)** | 51.12 ± 4.39 (A) | 51.12 ± 5.78 (A) |
|  | **Ps.S3D (mm²)** | 2252 ± 114 (A) | 2302 ± 164 (A) |

| **Table S4. Comparison of Pooled Lumbar Vertebrae (L4 - L6) qCT Parameters from Week 8 to Week 21.** This period during which animals were not given GC showed partial bone recovery with TMD.BV and BMD.Me increasing and BV and TV modestly rising, while BS/TV and BS/BV decreased indicating reduced surface irregularity. However, values at Week 21 were found to not have fully returned to baseline, suggesting incomplete reversal of osteoporosis. Values are presented as mean ± standard deviation where, across a parameter (*i.e.*, row), groups that possess different letters have statistically significant difference in mean (p ≤ 0.05) whereas those that possess the same letter have similar means (p > 0.05). N ≥ 12 (total vertebrae assessed per group). | | | | |
| --- | --- | --- | --- | --- |
|  | **Week 8** | **Week 13** | **Week 16** | **Week 21** |
| **TMD.BV (mgHA·cm⁻³)** | 479.2 ± 38.50 (A) | 528.2 ± 29.25 (B) | 569.6 ± 33.51 (C) | 609.2 ± 26.75 (D) |
| **TV (mm³)** | 2411 ± 228 (A) | 2345 ± 159 (A) | 2441 ± 169 (AB) | 2595 ± 144 (B) |
| **BV (mm³)** | 2203 ± 224 (AB) | 2141 ± 136 (A) | 2249 ± 155 (B) | 2396 ± 131 (C) |
| **BS (mm²)** | 1351 ± 743 (A) | 1459 ± 364 (A) | 1366 ± 317 (A) | 1106 ± 242 (A) |
| **BV/TV** | 0.914 ± 0.041 (A) | 0.913 ± 0.022 (A) | 0.922 ± 0.017 (A) | 0.924 ± 0.010 (A) |
| **BS/TV (mm⁻¹)** | 0.567 ± 0.330 (AB) | 0.624 ± 0.159 (A) | 0.567 ± 0.154 (AB) | 0.429 ± 0.098 (B) |
| **BS/BV (mm⁻¹)** | 0.636 ± 0.390 (AB) | 0.686 ± 0.183 (A) | 0.616 ± 0.171 (AB) | 0.464 ± 0.107 (B) |
| **BMD.Me (mgHA·cm⁻³)** | 262.4 ± 85.9 (A) | 317.1 ± 97.1 (AB) | 345.7 ± 105.8 (B) | 268.1 ± 52.8 (AB) |
| **TMD.Tb (mgHA·cm⁻³)** | 380.6 ± 78.7 (A) | 445.8 ± 80.1 (B) | 487.2 ± 77.7 (B) | 448.2 ± 42.0 (AB) |
| **Tb.Th (mm)** | 1.573 ± 0.266 (A) | 1.438 ± 0.180 (B) | 1.435 ± 0.119 (B) | 1.426 ± 0.115 (AB) |
| **Conn.D (mm⁻³)** | 0.147 ± 0.176 (A) | 0.334 ± 0.222 (B) | 0.287 ± 0.232 (AB) | 0.422 ± 0.244 (B) |
| **Anisotropy (SVD)** | 0.690 ± 0.186 (A) | 0.621 ± 0.112 (A) | 0.633 ± 0.111 (A) | 0.631 ± 0.113 (A) |
| **TMD.Co (mgHA·cm⁻³)** | 484.1 ± 56.2 (A) | 543.4 ± 36.7 (B) | 583.9 ± 33.5 (C) | 634.1 ± 25.3 (D) |
| **Ct.Ar (mm²)** | 66.98 ± 8.81 (A) | 64.90 ± 7.12 (A) | 66.32 ± 5.87 (A) | 68.50 ± 6.52 (A) |
| **Ct.Th (mm)** | 1.289 ± 0.533 (AB) | 1.127 ± 0.254 (B) | 1.210 ± 0.270 (AB) | 1.461 ± 0.197 (A) |
| **Tt.Ar (mm²)** | 66.98 ± 8.82 (A) | 64.90 ± 7.12 (A) | 66.32 ± 5.87 (A) | 68.50 ± 6.52 (A) |
| **Ct.Po** | 0.272 ± 0.176 (A) | 0.268 ± 0.090 (A) | 0.244 ± 0.078 (AB) | 0.172 ± 0.031 (B) |
| **Ps.Pm (mm)** | 51.12 ± 5.78 (A) | 51.13 ± 4.57 (A) | 52.69 ± 3.96 (A) | 53.43 ± 4.19 (A) |
| **Ps.S3D (mm²)** | 2302 ± 164 (AB) | 2286 ± 108 (A) | 2377 ± 128 (BC) | 2475 ± 75 (C) |

| **Table S5. Comparison of Pooled Lumbar Vertebrae (L4 - L6) qCT Parameters Between Week 0, Week 21, Week 24, and Week 27.** After partial recovery by Week 21, renewed GC exposure lowered mineral density (*i.e.*, TMD.BV, BMD.Me, and TMD.Co) and BV/TV by Week 0, while BS, BS/TV, and BS/BV increased reflecting a return of osteoporosis. These results confirm the reproducibility of GC-induced bone loss. Values are presented as mean ± standard deviation where, across a parameter (*i.e.*, row), groups that possess different letters have statistically significant difference in mean (p ≤ 0.05) whereas those that possess the same letter have similar means (p > 0.05). N ≥ 12 (total vertebrae assessed per group). | | | | |
| --- | --- | --- | --- | --- |
|  | **Week 0** | **Week 21** | **Week 24** | **Week 27** |
| **TMD.BV (mgHA·cm⁻³)** | 629.1 ± 31.90 (A) | 609.2 ± 26.75 (AB) | 585.4 ± 38.68 (B) | 547.7 ± 37.63 (C) |
| **TV (mm³)** | 2344 ± 156 (A) | 2595 ± 144 (B) | 2570 ± 228 (B) | 2489 ± 257 (B) |
| **BV (mm³)** | 2184 ± 147 (A) | 2396 ± 131 (B) | 2349 ± 190 (B) | 2239 ± 217 (AB) |
| **BS (mm²)** | 1192 ± 326 (A) | 1106 ± 242 (A) | 1337 ± 296 (A) | 1574 ± 305 (B) |
| **BV/TV** | 0.932 ± 0.012 (A) | 0.924 ± 0.010 (AB) | 0.915 ± 0.017 (B) | 0.901 ± 0.023 (C) |
| **BS/TV (mm⁻¹)** | 0.511 ± 0.143 (A) | 0.429 ± 0.098 (A) | 0.531 ± 0.149 (A) | 0.642 ± 0.158 (B) |
| **BS/BV (mm⁻¹)** | 0.549 ± 0.157 (A) | 0.464 ± 0.107 (A) | 0.580 ± 0.163 (A) | 0.715 ± 0.184 (B) |
| **BMD.Me (mgHA·cm⁻³)** | 357.5 ± 79.9 (A) | 268.1 ± 52.8 (B) | 311.1 ± 100.8 (AB) | 308.6 ± 102.7 (AB) |
| **TMD.Tb (mgHA·cm⁻³)** | 502.0 ± 52.1 (A) | 448.2 ± 42.0 (A) | 474.1 ± 70.9 (A) | 461.4 ± 82.0 (A) |
| **Tb.Th (mm)** | 1.537 ± 0.106 (A) | 1.426 ± 0.115 (AB) | 1.402 ± 0.156 (B) | 1.304 ± 0.150 (C) |
| **Conn.D (mm⁻³)** | 0.168 ± 0.137 (A) | 0.422 ± 0.244 (AB) | 0.379 ± 0.295 (B) | 0.465 ± 0.322 (B) |
| **Anisotropy (SVD)** | 0.679 ± 0.122 (A) | 0.631 ± 0.113 (AB) | 0.587 ± 0.099 (B) | 0.611 ± 0.102 (AB) |
| **TMD.Co (mgHA·cm⁻³)** | 656.5 ± 37.8 (A) | 634.1 ± 25.3 (AB) | 605.9 ± 41.1 (B) | 562.6 ± 46.1 (C) |
| **Ct.Ar (mm²)** | 65.16 ± 6.63 (A) | 68.50 ± 6.52 (A) | 65.71 ± 6.71 (A) | 63.88 ± 7.83 (A) |
| **Ct.Th (mm)** | 1.295 ± 0.274 (A) | 1.461 ± 0.197 (A) | 1.266 ± 0.246 (A) | 1.116 ± 0.248 (B) |
| **Tt.Ar (mm²)** | 65.16 ± 6.64 (A) | 68.50 ± 6.52 (A) | 65.71 ± 6.71 (A) | 63.89 ± 7.84 (A) |
| **Ct.Po** | 0.218 ± 0.068 (A) | 0.172 ± 0.031 (A) | 0.225 ± 0.064 (A) | 0.274 ± 0.091 (B) |
| **Ps.Pm (mm)** | 51.12 ± 4.39 (A) | 53.43 ± 4.19 (A) | 52.41 ± 3.92 (A) | 51.67 ± 4.77 (A) |
| **Ps.S3D (mm²)** | 2252 ± 114 (A) | 2475 ± 75 (B) | 2503 ± 140 (B) | 2467 ± 240 (B) |

| **Table S6. Post-VA Lumbar Comparison of qCT Parameters.** An overview of post-VA (Week 28) general qCT parameters for L4 - L6 with values is presented as mean ± standard deviation. To evaluate only bone parameters, PMMA volume was excluded prior to analysis. Across a parameter (*i.e.*, row), groups that possess different letters have statistically significant difference in mean (p ≤ 0.05) whereas those that possess the same letter have similar means (p > 0.05). N ≥ 5 (total vertebrae assessed per group). | | | |
| --- | --- | --- | --- |
|  | **L4** | **L5** | **L6** |
| **PMMA Volume (mm³)** | 37.47 ± 9.44 (A) | 31.11 ± 10.28 (A) | — |
| **TMD.BV (mgHA·cm⁻³)** | 551.9 ± 39.4 (A) | 557.0 ± 36.9 (A) | 567.7 ± 37.2 (A) |
| **TV (mm³)** | 2376 ± 204 (A) | 2553 ± 182 (A) | 2503 ± 222 (A) |
| **BV (mm³)** | 2167 ± 182 (A) | 2338 ± 178 (A) | 2324 ± 212 (A) |
| **BS (mm²)** | 1408 ± 323 (A) | 1322 ± 166 (A) | 1433 ± 285 (A) |
| **BV/TV** | 0.912 ± 0.014 (A) | 0.916 ± 0.017 (A) | 0.928 ± 0.010 (A) |
| **BS/TV (mm⁻¹)** | 0.599 ± 0.155 (A) | 0.524 ± 0.105 (A) | 0.583 ± 0.156 (A) |
| **BS/BV (mm⁻¹)** | 0.657 ± 0.171 (A) | 0.572 ± 0.112 (A) | 0.629 ± 0.169 (A) |
| **BMD.Me (mgHA·cm⁻³)** | 337.4 ± 113.0 (A) | 296.1 ± 112.4 (A) | 318.3 ± 102.3 (A) |
| **TMD.Tb (mgHA·cm⁻³)** | 492.9 ± 89.7 (A) | 455.5 ± 96.8 (A) | 445.9 ± 95.7 (A) |
| **Tb.Th (mm)** | 1.221 ± 0.170 (A) | 1.258 ± 0.255 (A) | 1.408 ± 0.118 (A) |
| **Conn.D (mm⁻³)** | 0.321 ± 0.194 (A) | 0.536 ± 0.383 (A) | 0.314 ± 0.180 (A) |
| **Anisotropy (SVD)** | 0.609 ± 0.104 (A) | 0.626 ± 0.040 (A) | 0.627 ± 0.094 (A) |
| **TMD.Co (mgHA·cm⁻³)** | 563.1 ± 31.2 (A) | 572.3 ± 29.9 (A) | 596.1 ± 34.9 (A) |
| **Ct.Ar (mm²)** | 60.84 ± 6.37 (A) | 64.99 ± 6.24 (A) | 70.05 ± 8.76 (A) |
| **Ct.Th (mm)** | 1.153 ± 0.255 (A) | 1.283 ± 0.192 (A) | 1.198 ± 0.259 (A) |
| **Tt.Ar (mm²)** | 60.84 ± 6.37 (A) | 64.99 ± 6.24 (A) | 70.05 ± 8.76 (A) |
| **Ct.Po** | 0.265 ± 0.073 (A) | 0.219 ± 0.055 (A) | 0.235 ± 0.072 (A) |
| **Ps.Pm (mm)** | 49.76 ± 3.09 (A) | 52.39 ± 3.41 (A) | 55.42 ± 4.67 (A) |
| **Ps.S3D (mm²)** | 2392 ± 113 (A) | 2520 ± 119 (A) | 2442 ± 136 (A) |

| **Table S7. Comparison of PMMA Injected L4 - L5 qCT Parameters Between Pre-VA (Week 27) and Post-VA (Week 28).** Values are presented as mean ± standard deviation where across a parameter (*i.e.*, row), groups that possess different letters have statistically significant difference in mean (p ≤ 0.05) whereas those that possess the same letter have similar means (p > 0.05). The “*” indicates p = 0.0568. N ≥ 10 (total vertebrae assessed per group). | | | |
| --- | --- | --- | --- |
|  |  | **Pre-VA**  **(Week 27)** | **Post-VA**  **(Week 28)** |
|  | **TMD.BV (mgHA·cm⁻³)** | 537.4 ± 35.7 (A) | 553.7 ± 36.6 (A) |
|  | **TV (mm³)** | 2476 ± 267 (A) | 2489 ± 215 (A) |
|  | **BV (mm³)** | 2213 ± 221 (A) | 2261 ± 196 (A) |
|  | **BS (mm²)** | 1609 ± 320 (A) | 1365 ± 232 (B) |
|  | **BV/TV** | 0.895 ± 0.023 (A)* | 0.909 ± 0.015 (A)* |
|  | **BS/TV (mm⁻¹)** | 0.660 ± 0.162 (A) | 0.555 ± 0.123 (B) |
|  | **BS/BV (mm⁻¹)** | 0.740 ± 0.191 (A) | 0.611 ± 0.136 (B) |
|  | **BMD.Me (mgHA·cm⁻³)** | 310.0 ± 102.9 (A) | 301.6 ± 106.4 (A) |
|  | **TMD.Tb (mgHA·cm⁻³)** | 461.9 ± 84.0 (A) | 468.8 ± 88.6 (A) |
|  | **Tb.Th (mm)** | 1.283 ± 0.154 (A) | 1.190 ± 0.183 (A) |
|  | **Conn.D (mm⁻³)** | 0.469 ± 0.347 (A) | 0.519 ± 0.326 (A) |
|  | **Anisotropy (SVD)** | 0.605 ± 0.106 (A) | 0.627 ± 0.076 (A) |
|  | **TMD.Co (mgHA·cm⁻³)** | 549.2 ± 43.5 (A) | 567.1 ± 29.6 (A) |
|  | **Ct.Ar (mm²)** | 61.36 ± 6.89 (A) | 63.83 ± 6.56 (A) |
|  | **Ct.Th (mm)** | 1.093 ± 0.254 (A) | 1.238 ± 0.206 (A) |
|  | **Tt.Ar (mm²)** | 61.37 ± 6.91 (A) | 63.83 ± 6.56 (A) |
|  | **Ct.Po** | 0.284 ± 0.094 (A) | 0.238 ± 0.060 (A) |
|  | **Ps.Pm (mm)** | 50.07 ± 4.35 (A) | 51.44 ± 3.45 (A) |
|  | **Ps.S3D (mm²)** | 2471 ± 271 (A) | 2464 ± 131 (A) |

| **Table S8. Comparison of Non-Injected L6 qCT Parameters Between Pre-VA (Week 27) and Post-VA (Week 28).** Values are presented as mean ± standard deviation where, across a parameter (*i.e.*, row), groups that possess different letters have statistically significant difference in mean (p ≤ 0.05) whereas those that possess the same letter have similar means (p > 0.05). N ≥ 5 (total vertebrae assessed per group). | | | |
| --- | --- | --- | --- |
|  |  | **Pre-VA**  **(Week 27)** | **Post-VA**  **(Week 28)** |
|  | **TMD.BV (mgHA·cm⁻³)** | 568.3 ± 33.5 (A) | 563.8 ± 35.5 (A) |
|  | **TV (mm³)** | 2514 ± 242 (A) | 2563 ± 204 (A) |
|  | **BV (mm³)** | 2292 ± 205 (A) | 2366 ± 195 (A) |
|  | **BS (mm²)** | 1505 ± 270 (A) | 1388 ± 271 (A) |
|  | **BV/TV** | 0.912 ± 0.017 (A) | 0.923 ± 0.013 (A) |
|  | **BS/TV (mm⁻¹)** | 0.608 ± 0.148 (A) | 0.550 ± 0.146 (A) |
|  | **BS/BV (mm⁻¹)** | 0.667 ± 0.164 (A) | 0.596 ± 0.159 (A) |
|  | **BMD.Me (mgHA·cm⁻³)** | 305.7 ± 105.4 (A) | 284.3 ± 98.9 (A) |
|  | **TMD.Tb (mgHA·cm⁻³)** | 460.3 ± 80.4 (A) | 422.8 ± 91.5 (A) |
|  | **Tb.Th (mm)** | 1.347 ± 0.135 (A) | 1.370 ± 0.101 (A) |
|  | **Conn.D (mm⁻³)** | 0.455 ± 0.275 (A) | 0.385 ± 0.207 (A) |
|  | **Anisotropy (SVD)** | 0.624 ± 0.094 (A) | 0.657 ± 0.074 (A) |
|  | **TMD.Co (mgHA·cm⁻³)** | 589.4 ± 39.8 (A) | 592.5 ± 32.8 (A) |
|  | **Ct.Ar (mm²)** | 68.92 ± 7.29 (A) | 71.50 ± 8.62 (A) |
|  | **Ct.Th (mm)** | 1.161 ± 0.236 (A) | 1.256 ± 0.237 (A) |
|  | **Tt.Ar (mm²)** | 68.93 ± 7.30 (A) | 71.51 ± 8.62 (A) |
|  | **Ct.Po** | 0.256 ± 0.084 (A) | 0.220 ± 0.066 (A) |
|  | **Ps.Pm (mm)** | 54.87 ± 3.97 (A) | 56.34 ± 4.66 (A) |
|  | **Ps.S3D (mm²)** | 2458 ± 167 (A) | 2488 ± 120 (A) |
